# Diffusion-Informed Spatial Smoothing of fMRI Data in White Matter Using Spectral Graph Filters

**DOI:** 10.1101/2020.10.25.353920

**Authors:** David Abramian, Martin Larsson, Anders Eklund, Iman Aganj, Carl-Fredrik Westin, Hamid Behjat

## Abstract

Brain activation mapping using functional magnetic resonance imaging (fMRI) has been extensively studied in brain gray matter (GM), whereas in large disregarded for probing white matter (WM). This unbalanced treatment has been in part due to controversies in relation to the nature of the blood oxygenation level-dependent (BOLD) contrast in WM and its detachability. However, an accumulating body of studies has provided solid evidence of the functional significance of the BOLD signal in WM and has revealed that it exhibits anisotropic spatio-temporal correlations and structure-specific fluctuations concomitant with those of the cortical BOLD signal. In this work, we present an anisotropic spatial filtering scheme for smoothing fMRI data in WM that accounts for known spatial constraints on the BOLD signal in WM. In particular, the spatial correlation structure of the BOLD signal in WM is highly anisotropic and closely linked to local axonal structure in terms of shape and orientation, suggesting that isotropic Gaussian filters conventionally used for smoothing fMRI data are inadequate for denoising the BOLD signal in WM. The fundamental element in the proposed method is a graph-based description of WM that encodes the underlying anisotropy observed across WM, derived from diffusion-weighted MRI data. Based on this representation, and leveraging graph signal processing principles, we design subject-specific spatial filters that adapt to a subject’s unique WM structure at each position in the WM that they are applied at. We use the proposed filters to spatially smooth fMRI data in WM, as an alternative to the conventional practice of using isotropic Gaussian filters. We test the proposed filtering approach on two sets of simulated phantoms, showcasing its greater sensitivity and specificity for the detection of slender anisotropic activations, compared to that achieved with isotropic Gaussian filters. We also present WM activation mapping results on the Human Connectome Project’s 100-unrelated subject dataset, across seven functional tasks, showing that the proposed method enables the detection of streamline-like activations within axonal bundles.

## 1. Introduction

To date, reports on task-based functional magnetic resonance imaging (fMRI) activation mapping and resting-state functional connectivity have been overwhelmingly restricted to the gray matter (GM), whereas white matter (WM) functional data have been largely ignored or treated as a nuisance regressor. Such unbalanced treatment of fMRI data within GM and WM, due in part to controversies in relation to the source of the BOLD signal in WM, has led to a systematic underreporting of BOLD-related activity in WM (Mazerolle et al., 2019; Gawryluk et al., 2014).

Despite past controversies, evidence provided by an increasing body of recent studies, see e.g. Grajauskas et al. (2019) and Gore et al. (2019) and references therein, has led to more widespread acceptance of the detectability and functional relevance of the BOLD signal in WM. For example, Ding et al. (2013) showed that resting-state BOLD signals in WM exhibit structure-specific temporal correlations along WM tracts, which coincide with fiber patterns revealed by diffusion tensor imaging (DTI), and which, under functional load, become more pronounced in functionally relevant structures (Ding et al., 2016). More specifically, Mishra et al. (2020) showed that varying experimental task parameters results in a coupled modulation of the BOLD signal in the visual cortex and relevant WM tracts, corroborating past findings of simultaneous BOLD activations in structurally-connected regions of GM and WM (Mazerolle et al., 2010). More recently, it has been shown that functional neuroplasticity, as manifested by changes in the BOLD signal, can be detected in WM (Frizzell et al., 2020). Furthermore, a growing number of recent studies have shown that low frequency BOLD fluctuations can be used to estimate the dynamic functioning of fiber tracts (Gore et al., 2019), in both health (Marussich et al., 2017; Huang et al., 2018b; Li et al., 2020b) and disease (Jiang et al., 2019; Ji et al., 2019; Gao et al., 2020), providing a powerful means to study how information is transferred and integrated between functionally specialized cortices.

Due to the significantly lower vascularization density in WM compared to that in GM (Logothetis and Wandell, 2004; Jochimsen et al., 2010), the overall magnitude of the BOLD signal in WM is substantially lower than that in GM (Yarkoni et al., 2009), which has been reported to be as low as 10% of that observed in GM and modulated as a function of distance from the cortical layer (Li et al., 2019b). In addition to being weak, the BOLD signal in WM is affected by unique confounding factors, suggesting the need for WM-tailored acquisition and processing schemes. Broadly speaking, the BOLD contrast and its detection in WM can potentially be enhanced in three ways: i) development and use of MRI sequences optimal for fMRI of WM (e.g. increased T2-weighting (Gawryluk et al., 2009) or tailored field strengths (Mazerolle et al., 2013)); ii) design of temporal models that account for the unique hemodynamic response function (HRF) in WM, which substantially differs from that in GM (Yarkoni et al., 2009; Fraser et al., 2012; Erdoğan et al., 2016)); and iii) design of spatial models that account for the unique spatial features of the BOLD contrast in WM, which is highly anisotropic (Ding et al., 2013, 2016). This paper focuses on the third category, presenting the case for the importance of spatial filter design when handling fMRI data in WM, particularly in relation to the inherent differences between the spatial profiles of BOLD signal in WM relative to those in GM.

### 1.1 Spatial smoothing tailored to fMRI data in white matter

Typical fMRI analysis pipelines rely on the assumption that the BOLD signal exhibits isotropic spatial profiles at focal activated regions (Carp, 2012). Isotropic Gaussian kernels applied to functional data, which is a staple of conventional fMRI analysis, is only justified under this assumption, and generally trades spatial specificity for increased sensitivity. In particular, by virtue of the matched filter argument, spatial filters are optimal only for detecting activations that conform to the size and shape of the filter kernel, and can otherwise result in loss of information regarding the spatial extent and shape of activation areas (Geissler et al., 2005; Mikl et al., 2008), obliterating all non-smooth singularities in the data.

In order to improve on the sensitivity-specificity trade-off afforded by conventional isotropic spatial smoothing, multiple smoothing methods that adapt to local spatial image features have been proposed. These include steerable filters (Knutsson et al., 1983), which enable directionally-adaptive spatial smoothing (Friman et al., 2003; Eklund et al., 2011; Zhuang et al., 2017; Abramian et al., 2020b), wavelet transforms (Mallat, 1989; Bullmore et al., 2004), which try to strike a balance between localization in space and frequency domain (Ruttimann et al., 1998; Van De Ville et al., 2004; Breakspear et al., 2006), and non-linear filters (e.g. bilateral filters) that locally adapt to various features of adjacent voxels (Smith and Brady, 1997; Rydell et al., 2008; Lohmann et al., 2018). While such methods have been successfully applied to GM, their adaptive properties rely on the spatial features manifested by the BOLD contrast. Given that this contrast is substantially reduced in WM, the effectiveness of these methods would likely be reduced when applied to fMRI data in WM.

Rather than adapting the smoothing operation to features present in the BOLD contrast, alternative adaptive smoothing approaches can be leveraged that incorporate information from the *domain* on which the data reside, typically provided by complementary anatomical images. One common approach is cortical surface smoothing, which has shown to provide increased sensitivity and specificity (Jo et al., 2007; Coalson et al., 2018). Such methods have also been used to formulate smoothing approaches that respect tissue boundaries (Behjat et al., 2019), preventing artifacts resulting from the mixing of signals from adjacent but differing tissue types during filtering. In both of these scenarios the anatomical information is provided by T1-weighted images.

An important distinguishing feature of the BOLD signal in WM is that it exhibits a spatial correlation structure grossly consistent with the directions of water diffusion, as measured by DTI (Ding et al., 2013), which is present during rest and becomes more pronounced under functional loading (Wu et al., 2017; Ding et al., 2018). The anatomical basis for this observation can be that up to half of the blood volume in WM resides in vessels that run in parallel to WM tracts (Doucette et al., 2019). As a consequence, conventional isotropic Gaussian filters may prove especially unsuited for the task of increasing the SNR of the BOLD signal in the highly anisotropic WM domain. Filtering methods adaptive to features of the BOLD signal may prove more effective, but the low BOLD contrast manifested in WM will potentially limit their usefulness. On the other hand, the strong anatomical dependence in the correlation structure of the BOLD signal in WM suggests that domain-informed smoothing methods can be particularly beneficial. Such methods can rely on T1-weighted images as well as diffusion-weighted MRI (DW-MRI) to adapt the filtering to the morphology and the axonal microstructure of WM, respectively. This paper presents the design and validation of such a filtering scheme.

### 1.2. Structure-informed processing of fMRI data through GSP

In the past five years, an increasing number of studies have showcased the use of principles from the recently emerged field of graph signal processing (GSP) within neuroimaging, in particular, in proposing intuitive methodologies for structure-informed processing of fMRI data. The fundamental idea in GSP is to analyze data recorded at a discrete set of positions in such way that the underlying structural relationship between those positions is accounted for, wherein this underlying structure can be represented in the form of a graph, i.e., a structure consisting of a set of vertices and edges. We refer the reader to Shuman et al. (2013) for an introduction to GSP and to Ortega et al. (2018) and Stanković et al. (2020) for an overview of recent developments, challenges, and applications.

An increasing number of studies have proposed the use of region of interest (ROI) based structural connectomes (Sporns et al., 2005), derived from tractography data, as underlying backbones for interpreting fMRI data (Atasoy et al., 2016; Abdelnour et al., 2018; Huang et al., 2018a). When structural connectomes are interpreted as graphs, a number of their Laplacian eigenvectors manifest spatial patterns that are reminiscent of well-established functional networks, as shown by Atasoy et al. (2016). Under this framework, methods have been proposed for spatio-temporal deconvolution of fMRI data (Bolton et al., 2019), quantification of the coupling strength of resting-state fMRI data with underlying structure (Medaglia et al., 2018; Preti and Van De Ville, 2019), implementation of neural field models (Aqil et al., 2020), prediction of brain disorders (Itani and Thanou, 2020) or behaviorally relevant scores (Bolton and van De Ville, 2020), and for characterization of functional connectivity dynamics in health (Huang et al., 2018b), and its changes, for instance, due to concussion (Sihag et al., 2020), and under hallucinogenic drugs (Atasoy et al., 2017).

As alternatives to macro-scale ROI-based graphs, a number of voxel-wise brain graph designs have been proposed for analysis of fMRI data. Graphs encoding GM morphology have been proposed for enhanced activation mapping in GM, for both group-level (Behjat et al., 2015) and subject-level (Behjat et al., 2013, 2014) analyses, and for discriminative characterization of fMRI data across functional tasks (Behjat and Larsson, 2020). A closely related work to that presented here is by Tarun et al. (2020), in which DW-MRI data were used to encode the WM fiber structure, for the task of visualizing WM fiber pathways based on the functional activity observed at the cortical layer.

### 1.3. Aim and overview

To the best of our knowledge, no method has to date been presented to specifically account for the spatial features of the BOLD contrast in WM when it comes to spatial processing of fMRI data. The main objective of this work is to present the case for the importance of spatial filter design when handling fMRI data in WM, particularly, in relation to the inherent difference between the spatial profiles of BOLD signal in WM relative to those in GM.

In this paper, we develop an adaptive spatial smoothing method tailored to the processing of fMRI data in WM. Using diffusion orientation distribution functions (ODF) obtained from high angular resolution diffusion imaging (HARDI) data, we construct subject-specific voxel-wise WM graphs. A spectral heat kernel filter is then defined on the spectrum of the resulting graphs, and implemented in a computationally efficient way for the task of fMRI data filtering, using principles from GSP. When instantiated at any position within the WM, the proposed filters adapt to the local axonal orientation, becoming consistent with the spatial correlation structure of the BOLD signal in WM.

The remainder of this paper is organized as follows: in Section 2, we review relevant GSP principles and describe our proposed graph and filter designs, as well as the construction of phantoms. In Section 3, we examine the smoothing filters produced by the proposed design and evaluate their performance on phantoms of two types and on real task fMRI data. We conclude the paper in Section 4 with a discussion on design considerations, limitations and future work.

## 2. Materials and Methods

### 2.1. Data and preprocessing

Data used in the preparation of this work were obtained from the WU-Minn Human Connectome Project (HCP) (Van Essen et al., 2013) database^1^. We use the 100 unrelated adult subject sub-group (54% female, mean age = 29.11 ± 3.67, age range = 22-36), which we denote as the HCP100 subject set. Five of the subjects were excluded due to incomplete WM coverage of the DW-MRI data, leaving a total of 95 subjects. The HCP data acquisition study was approved by the Washington University Institutional Review Board and informed consent was obtained from all subjects. We used the minimally preprocessed structural, task fMRI, and DW-MRI data. Task fMRI data for each subject consist of 1940 time frames across seven functional tasks: Emotion, Gambling, Language, Motor, Relational, Social, and Working Memory, comprising 23 experimental conditions in total. The method proposed in this paper heavily relies on the accurate co-registration between the structural and functional data, as provided by the minimally processed HCP data. The imaging parameters and image preprocessing steps have been thoroughly described by Glasser et al. (2013). All data processing in this work was done using the MATLAB software and the SPM12 toolbox^2^. Diffusion ODFs were generated using the method presented by Yeh et al. (2010) and implemented in the DSI Studio software packagee^3^.

The HCP preprocessed data are provided in a mixture of three spatial resolutions within two neurological spaces (ACPC, i.e., native subject space, and MNI): 0.7 mm isotropic ACPC for the structural data, 1.25 mm isotropic ACPC for the DW-MRI data, and 2 mm isotropic MNI for the fMRI data. A fundamental necessity for the proposed methodology is to reconcile the three datasets into a single set of working parameters. However, the resampling process and the nonlinear conversion between ACPC and MNI spaces have the potential of negatively affecting the data quality. The number of voxels is also a relevant parameter, as it determines to a great extent the memory usage and computation time of the various processing steps. Given the importance of axonal orientation information to the proposed method, we prioritized minimizing the manipulations applied to the DW-MRI data.

Based on these considerations, we chose the ACPC space at the resolution of the diffusion data, i.e., 1.25 mm isotropic, as the working space. As such, the HCP preprocessed fMRI volumes were warped back into ACPC space and upsampled to the voxel resolution of the diffusion data. This mapping was done by leveraging the `mni2acpc.nii` displacement maps provided with the HCP preprocessed data, using first order splines as the basis for interpolation. In addition, the segmentation volume `aparc+aseg.nii`, computed via FreeSurfer (Fischl, 2012) and provided with the HCP data, was downsampled to the working resolution, from which voxels associated to WM were extracted.

### 2.2. GSP preliminaries

The fundamental idea in GSP is the application of signal processing procedures to data residing on the vertices of a graph, wherein the graph defines the underlying irregular domain of the data. Let *𝒢*= (*𝒱, ε*, **A**) denote an undirected, connected, weighted graph, defined by a vertex set *𝒱* of size *N*_*g*_, denoting the size of the graph, an edge set *ε* consisting of connecting pairs (*i, j*) of vertices, and a symmetric adjacency matrix **A** whose nonzero elements *a*_*i, j*_ represent the weight of edges (*i, j*) ∈ *ε*. Let *ℓ*^2^(*𝒢*) denote the Hilbert space of all square-integrable graph signals **f** : *𝒱* → ℝ defined on the vertex set *𝒱*. A graph signal **f** ∈ *ℓ*^2^ (*𝒢*) is in essence an *N*_*g*_ × 1 vector, whose *n*-th component represents the signal value at the *n*-th vertex of *𝒢*.

The graph spectral domain, analogous to the Euclidean Fourier domain, can be defined using a graph’s Laplacian matrix. In particular, the *normalized* Laplacian matrix *𝒢* of is defined as **L** = **I** − **D**−1/2**AD**−1/2, where **D** denotes the graph’s degree matrix, which is diagonal with elements defined as *d*_*i,i*_ = ∑*j a*_*i, j*_. Given that **L** is real, symmetric, diagonally dominant, and with non-negative diagonal entries, it is positive semi-definite; i.e., all its *N*_*g*_ eigenvalues are real and non-negative, and they are also no larger than 2 due to the normalization used in the definition of **L**. This set of eigen-values defines the spectrum of *𝒢* (Chung, 1997), denoted as 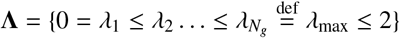 The associated eigen-vectors, denoted 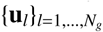, form an orthonormal basis spanning the *ℓ*^2^(𝒢) space.

In classical Fourier analysis, complex exponentials of varying frequencies are used to obtain spectral representations of signals, with larger frequencies corresponding to greater variability—per region or unit of time. It can be shown that, in the graph setting, the eigenvalues and eigenvectors of **L** fulfill a corresponding role to the frequencies and complex exponentials of the classical domain, respectively. In particular, larger eigenvalues of **L** are similarly associated to eigenvectors with greater spatial variability; we refer the interested reader to Appendix A for a more detailed presentation of this point. Given this analogy between the classical and graph settings, the eigen-vectors of **L** can be used to obtain spectral representations of graph signals. Specifically, a graph signal **f** can be transformed into a spectral representation through the use of the Laplacian eigenvectors as

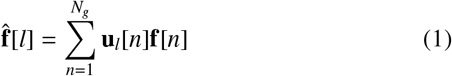

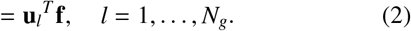

This spectral representation possesses a perfect reconstruction, that is, the signal can be recovered 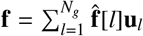.

In contrast to filters in classical signal processing, graph filters are *shift-variant*, adapting their shape to the underlying graph structure when localized at any given vertex. Consequently, individual filters defined in the spectral domain of a graph will become spatially-adaptive by the nature of GSP. This valuable property of graph filters enables the proposed methodology, but it also prevents the implementation of filtering operations as straightforward convolutions. Instead, in analogy to frequency-domain filtering in classical signal processing, graph signal filtering can be conveniently defined in the graph spectral domain. Given the spectral profile of a desired filter, *k*(λ) : [0, 2] → ℝ, a graph signal **f** can be filtered with *k*(λ) as

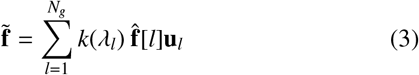

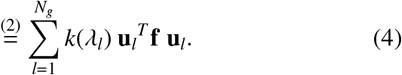

However, implementing (4) requires the Laplacian eigenvectors, i.e., a full diagonalization of **L**, which is impractical for large graphs, such as those presented in this work. An efficient alternative approach is to implement the filtering using a polynomial approximation of *k*(λ) (Hammond et al., 2011). We refer the interested reader to Appendix B for details on the implementation.

### 2.3. WM graph design

In order to take advantage of GSP tools, it is necessary to define graphs that encode relevant information in their vertices, edges, and weights. For the purpose of allowing diffusion-informed smoothing of the BOLD signal in WM, we require graphs capable of encoding the subject’s axonal microstructure. Filters defined on the spectral domain of such graphs will become locally adapted to this microstructure due to the shift-variant nature of graph filters.

We define a WM graph as a graph whose vertex set *𝒱* consists of all WM voxels, resulting in graphs with 240k ± 60k vertices on the HCP100 subject set. The graph’s edge *ε* set is defined on the basis of voxel adjacency, with pairs of vertices being connected to each other whenever their associated voxels are spatially neighboring. Two neighborhood definitions are considered, corresponding to cubic lattices of sizes 3 × 3 × 3 (henceforth 3-conn) and 5 × 5 × 5 (henceforth 5-conn), where the focal voxel is located in the center of the lattice. The 3-conn lattice specifies 26 voxels in the neighborhood of the focal voxel, whereas for the 5-conn lattice, voxels in the outer layer that fall in parallel to the voxels within the inner layer are excluded, resulting in 98 voxels in the neighborhood; see Figure 1.

**Figure 1:**
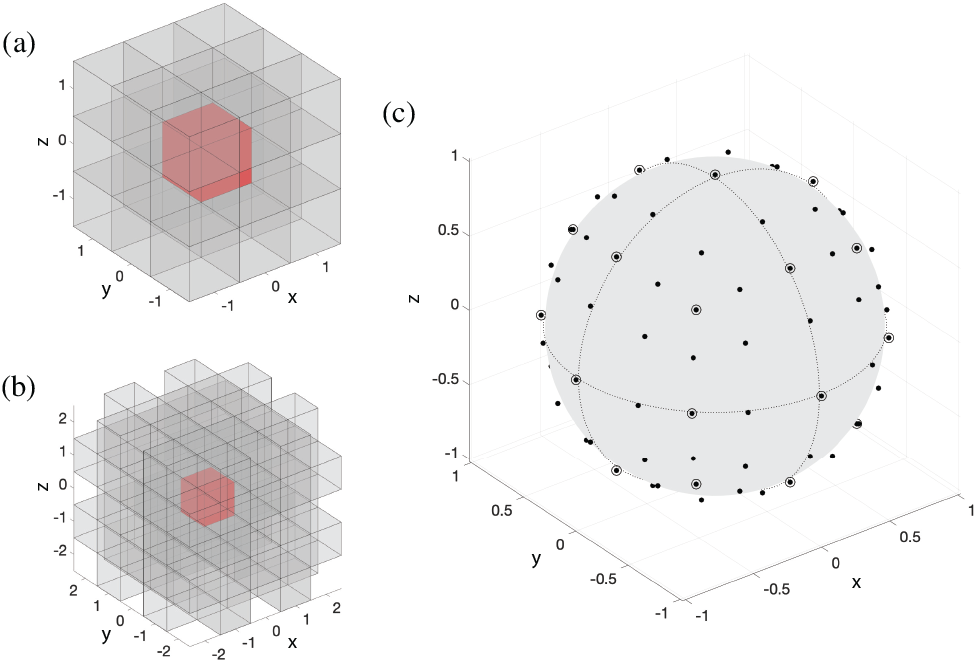
(a) 26 voxels within the 3 × 3 × 3 neighborhood (gray) used to define edges to the focal voxel (red). (b) 98 voxels within the 5 × 5 × 5 neighborhood (gray), used to define edges to the focal voxel (red). (c) Scattered dots on the unit sphere specify the 98 neighborhood directions encoded by the 5 × 5 × 5 voxel neighborhood. Circled dots represent the subset of 26 directions encoded by the 3 × 3 × 3 voxel neighborhood.

The encoding of axonal microstructure by the graph is principally achieved through the edge-weighting scheme, inspired by the work of Iturria-Medina et al. (2007). The weights provide a discretization of the diffusion ODF at each point, and include information on the coherence of diffusion orientation among neighboring voxels. Let 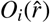 denote the ODF associated to voxel *v*_*i*_, with its coordinate origin at the voxel’s center, and with 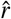 denoting the unit direction vector. Let 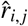 denote the unit vector pointing from the center of voxel *v*_*i*_ to the center of neighboring voxel *v*_*j*_. A discretization of the ODF along direction 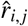 can be obtained as

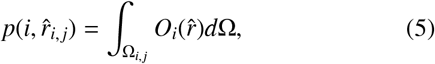

where Ω_*i,j*_ denotes the solid angle of 4π/26 (for 3-conn) or 4π/98 (for 5-conn) around 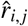 and *d*Ω denotes the infinitesimal solid angle element. This measure can be approximated by taking *N*_*t*_ samples of the ODF within the solid angle Ω_*i,j*_ as

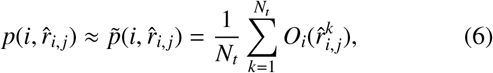

where 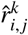 denotes the *k*-th sampling direction within Ω_*i,j*_. Details of the sampling process are given in Appendix C. Furthermore, we normalize this metric as

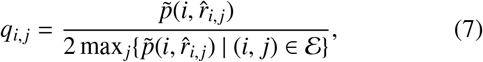

which bounds it in the [0, 0.5] range. The maximum value of 0.5 is reached if the ODF at *v*_*i*_ shows its maximal diffusion along 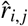, whereas otherwise *q*_*i,j*_ < 0.5.

The measure defined in (7) constitutes a normalized discretization of the diffusion ODF at voxel *v*_*i*_. However, it does not guarantee symmetry, i.e., generally *q*_*i,j*_ ≠ *q*_*j,i*_, which makes it unsuitable for the edge weights in an undirected graph. Nevertheless, we can obtain a symmetric weight by considering a bidirectional measure of diffusion given by

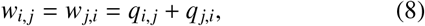

which is constrained to the [0, 1] range. Consequently, we define the graph’s edge weights as

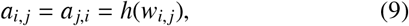

where *h*(·) : [0, 1] → [0, 1] is a tunable sigmoid function (Granlund and Knutsson, 1994) defined as

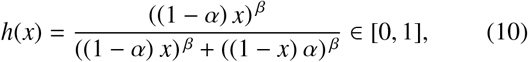

where parameters *α* ∈ (0, 1) and *β >* 0 control the threshold level and the steepness of the transition from 0 to 1, respectively; see Figure 2. Given that diffusion ODFs generally manifest non-zero magnitudes in all directions, with little contrast between directions of strong and weak diffusion, the thresholding step enables associating weights only to the main directions of diffusion, without the need to use sharpened ODFs as presented in our preliminary work (Abramian et al., 2020a). The choice of the sigmoid function over a Heaviside step ensures retaining a single connected structure in the graph; that is, any non-zero value is mapped to a non-zero value. In this work we use a fixed value of *β* = 50, but study the effect of varying the threshold point, in particular, for values *α* = 0.85, 0.9 and 0.95.

**Figure 2:**
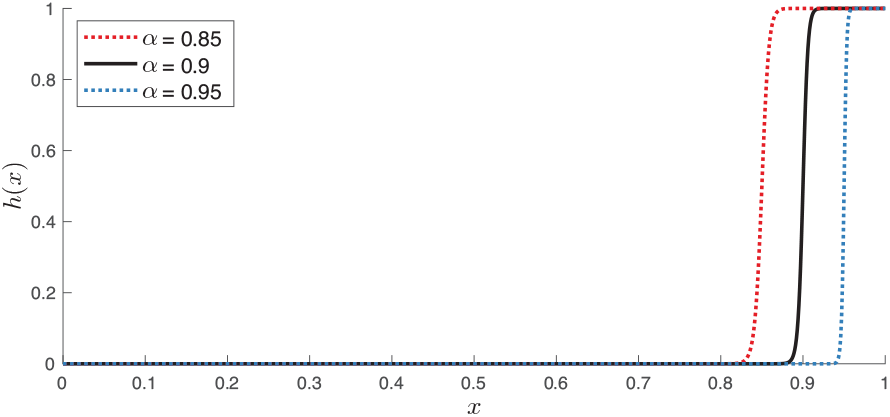
Sigmoid function used for thresholding edge weights, for three different values of *α* and a fixed value *β* = 50.

The expression for the edge weight between a pair of voxels (9) integrates information about the extent of diffusion along 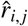 from both *v*_*i*_ and *v*_*j*_, amounting to a measure of orientational coherence of the diffusion ODFs at these voxels. In addition, the *α* parameter of the thresholding function provides added flexibility to this representation.

### 2.4. Spectral graph heat kernel filters

We design spatial smoothing filters with a heat kernel profile in the graph spectral domain, defined by

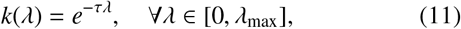

where *τ* is a free parameter determining the spatial extent of the filter. Figure 3 shows several realizations of the heat kernel over a range of *τ*. When instantiated in the vertex domain, such filters are roughly similar in shape to the Gaussian filters typically used for fMRI analysis; however, given the irregular domain represented by the graph, there is no direct equivalence between the two filters.

**Figure 3:**
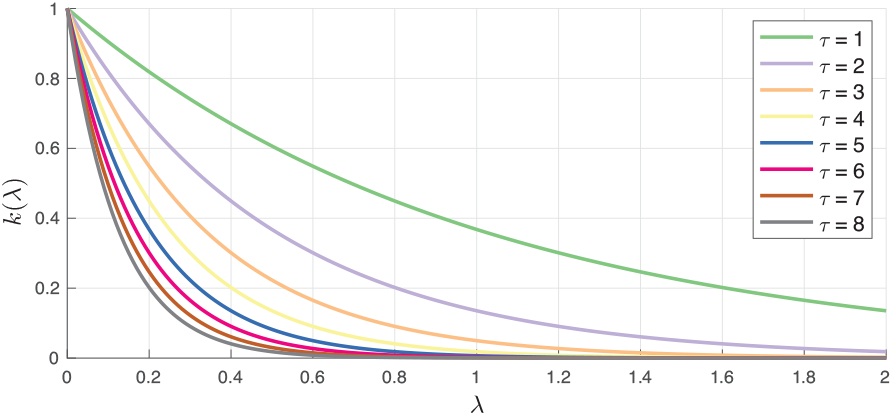
Spectral graph heat kernels, defined within the bounds of the spectrum of a normalized graph Laplacian matrix, i.e., [0, 2].

The filtering is implemented using the polynomial approximation scheme described in Appendix B. The polynomial order required to obtain a suitable approximation of the heat kernel varies depending on the choice of *τ*. For the range of *τ* investigated in this study, we used polynomial approximations of order 15, resulting in negligible approximation error in representing the filters.

### 2.5. Circular phantom construction

Due to the discrete nature of graphs, the set of orientations that can be perfectly captured by edges between voxels is limited by the neighborhood definition used. To evaluate the influence of angular resolution on denoising performance, we tested the 3-conn and 5-conn neighborhood definitions on a set of simulated circular phantoms of various orientations and radii. These phantoms aim to simulate a wide range of streamline orientations and curvatures, which could be encountered in practice.

Each phantom consisted of an activation profile in the shape of a circular streamline, accompanied by an ODF map oriented along its tangent, representing strong diffusion along the circle. The phantoms were constructed in 93 different orientations in 3D space, selected in a roughly uniform way by subdividing the faces of an icosahedron three times, and from the resulting polyhedron, selecting its subset of vertices that fall in the spherical sector of 0 ≤ *θ, ϕ* ≤ *π*/2; see Figure 4(a). Due to symmetries in the phantoms and the neighborhood definitions, this set of phantom orientations provides a relatively exhaustive sampling of the effects of streamline orientation on smoothing performance. Additionally, to study the effects of curvature, we created the phantoms with three different radii for each orientation: 10, 20, and 30 voxels at 1.25 mm isotropic resolution.

**Figure 4:**
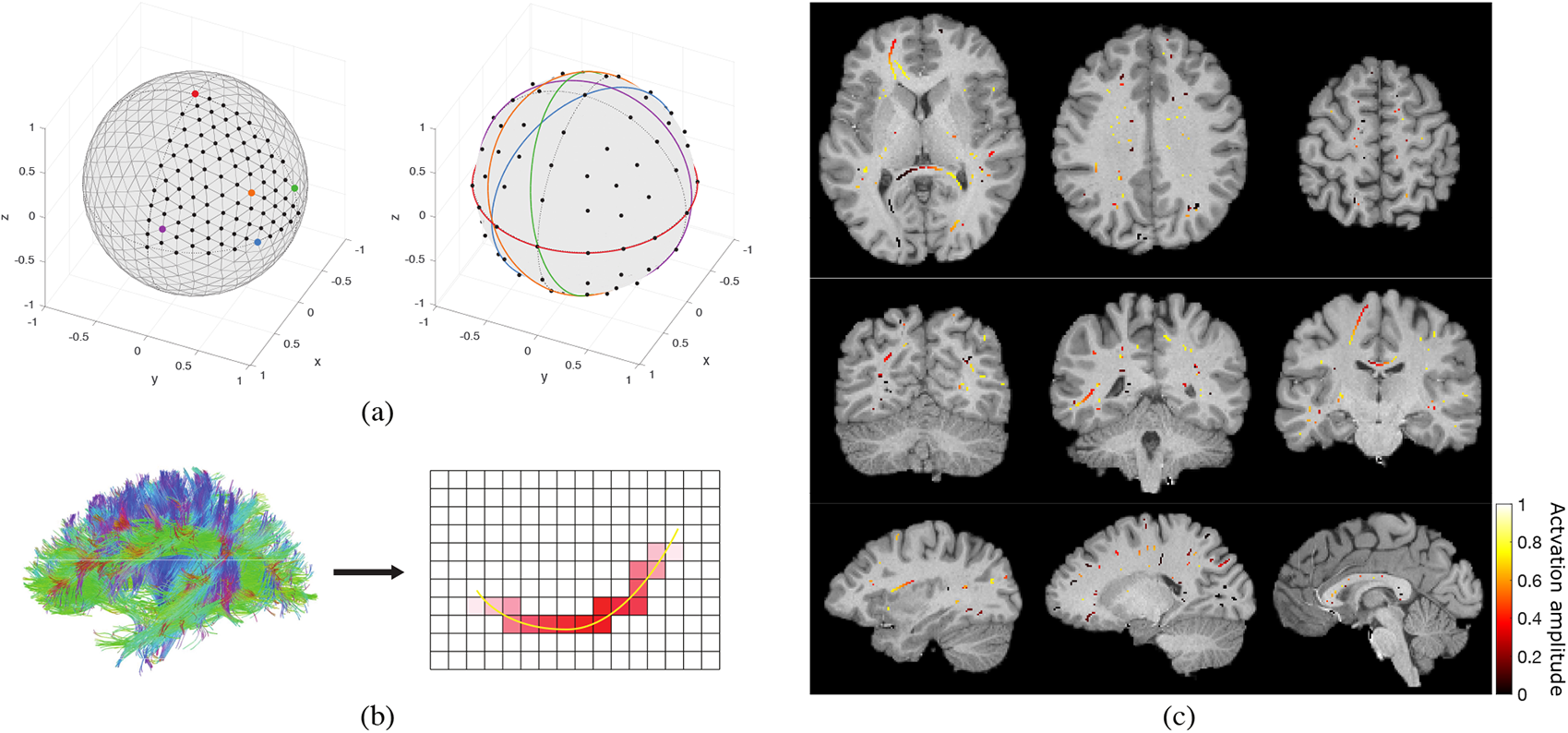
Phantom construction. (a) Circular phantom construction. Left: A subset of vertices of a 3-level subdivided icosahedron, 93 out of 642, were selected. Vectors pointing from the center of the sphere to these vertices constitute the normal vectors of the planes within which circular phantoms were realized. Right: Five representative unit circles with orientations corresponding to the vertices on the left of matching color. For example, the red circle falls within a plane that passes through the center of the sphere and has its normal vector pointing from the center of the sphere to the red point shown on the left. (b) Streamline-based phantom construction. A WM streamline constructed using tractography (shown in yellow) is randomly selected, a focal point along the streamline is randomly selected, and a diffused non-binary activation pattern is created around the focal point (shown in red). (c) Axial, coronal, and sagittal view of a representative streamline-based phantom with 100 streamline activations, overlaid on subject’s T1-weighted image.

### 2.6. Streamline-based phantom construction

Given that the correlation structure of the BOLD signal in WM is highly anisotropic and resemblant of the diffusion tensor (see Section 1.1), activation patterns in this tissue are likely to have elongated shapes which locally follow the direction of diffusion. To validate the performance of the proposed filtering scheme at detecting such activation patterns, we performed tests on a set of simulated semi-synthetic phantoms that simulate streamline-shaped activations. We denote the phantoms as *semi-synthetic*, as the spatial activation patterns were derived from real diffusion data from the HCP100 dataset. Each phantom consisted of a set of non-uniformly spread activation patterns diffusing along WM streamlines obtained through deterministic tractography of the HCP100 subject set; see Figures 4(b) and (c). Details of the construction of the phantoms are given in Appendix D.

Time-series versions of the streamline-based phantoms were also generated in order to evaluate the performance of the proposed method in the context of a typical fMRI general linear model (GLM) analysis. These were created by using each streamline-based phantom as the underlying ground-truth activity in a 100-volume fMRI time series, with a block design alternating 20 volume stretches of rest and activity in an off-on-off-on-off paradigm.

## 3. Results

We validated the performance of the proposed diffusion-informed spatial smoothing (DSS) method relative to isotropic Gaussian spatial smoothing (GSS) through a series of tests on synthetic phantoms—circular and streamline-based—and produced proof-of-concept results on real data from the HCP100 subject set.

### 3.1. Diffusion-informed filters

The adaptive properties of DSS filters are illustrated in Figure 5. The three filters shown were generated using identical parameters (α = 0.9, τ = 7), and differ only in the location within the WM where they were instantiated. The filters closely follow the local diffusion orientation in WM described by the diffusion ODFs. For highly anisotropic WM regions this results in slender and strongly oriented filters—see first two columns, whereas for regions of low anisotropy it results in filters that are more isotropic in shape. Particularly, at crossing fiber regions, DSS filters are not constrained to follow any single axonal pathway, and instead spatially extend along all directions of high diffusion—see third column. This avoids the uncertainty inherent in resolving the orientation of individual crossing fibers, while still resulting in more spatially-constrained filters than would be achieved with isotropic Gaussian filtering.

**Figure 5:**
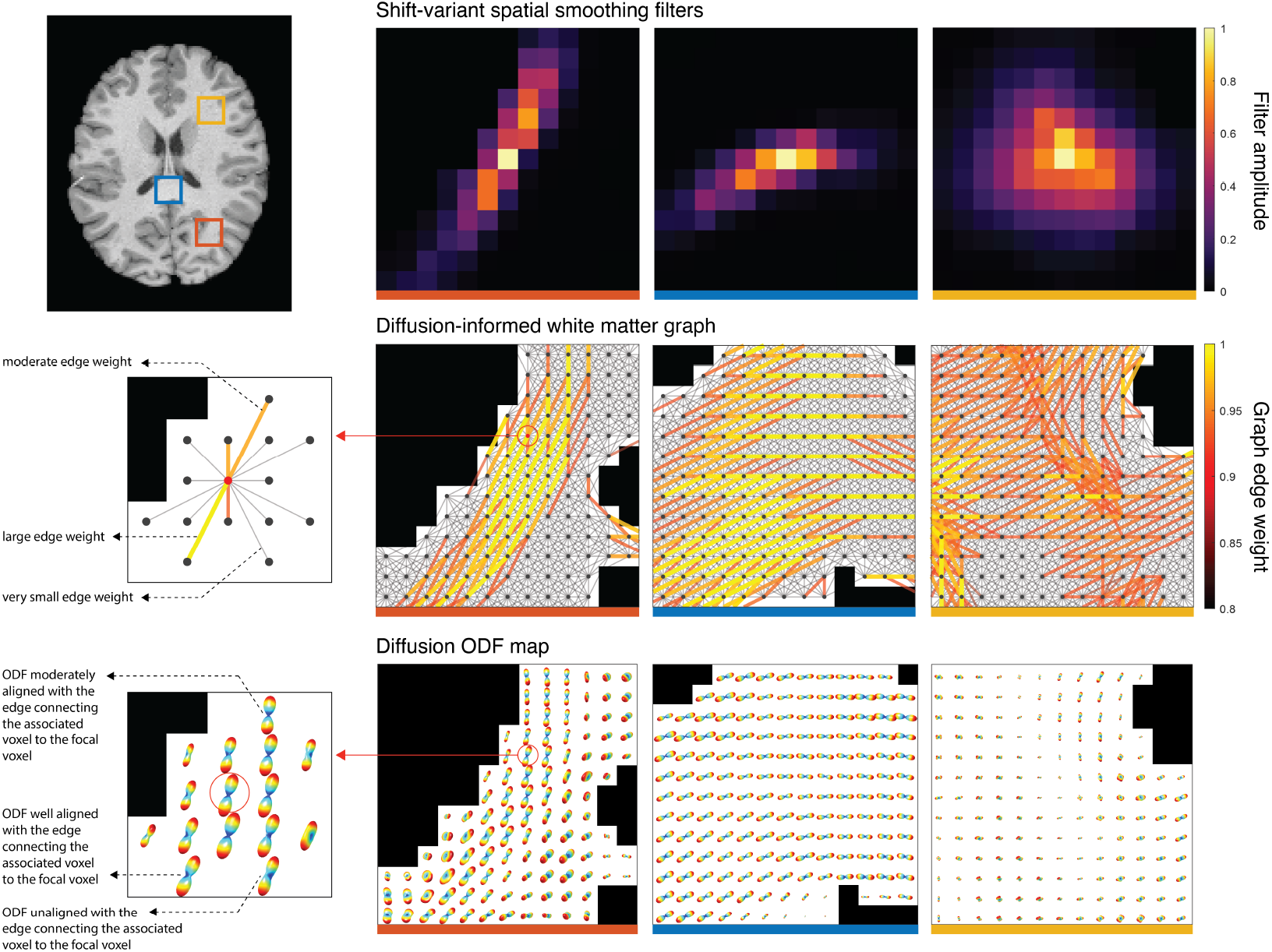
Generation of diffusion-informed smoothing filters. Diffusion ODFs (bottom row) serve as the basis for the creation of a WM graph (middle row). Every WM voxel corresponds to a vertex in the graph, with weighted connections to neighboring voxels (middle left). The edge weights are determined on the basis of coherence between the directions of diffusion and the orientation of the graph edges (bottom left). Using this WM graph definition, graph filters from a single spectral profile become adaptive to the local axonal microstructure when instantiated in different WM regions (top row). Note that both the edges connecting voxels and the graph filters extend in three dimensions, whereas their 2D axial intersection centered at the focal voxel are shown. Graph parameters: 5-conn neighborhood, α = 0.9, *β* = 50; filter parameters: τ = 7. Filters are shown normalized to the [0, 1] range. ODF interpolation and visualization were performed using the public CSA-ODF package^4^.

The shape of DSS filters can be controlled by setting the τ parameter of the graph spectral filter kernel (see (11)) and the α parameter of the weight thresholding function (see (10)). While the former controls the spatial extent of the filter in a manner akin to the full width at half maximum (FWHM) of isotropic Gaussian filters, the latter controls the minimum edge weights retained by the graph, which in turn, constrains filters to follow main directions of diffusion. Figure 6 presents a range of different filter shapes that can be achieved by varying these two parameters. High values of α result in very narrow, streamline-like filters that are highly constrained relative to the underlying diffusion map, whereas lower values result in less constrained filters. In particular, low enough values of α negate the diffusion-adaptive properties of DSS, with the resulting filters adapting solely to the morphology of the WM domain (see Supplementary Figure S1).

**Figure 6:**
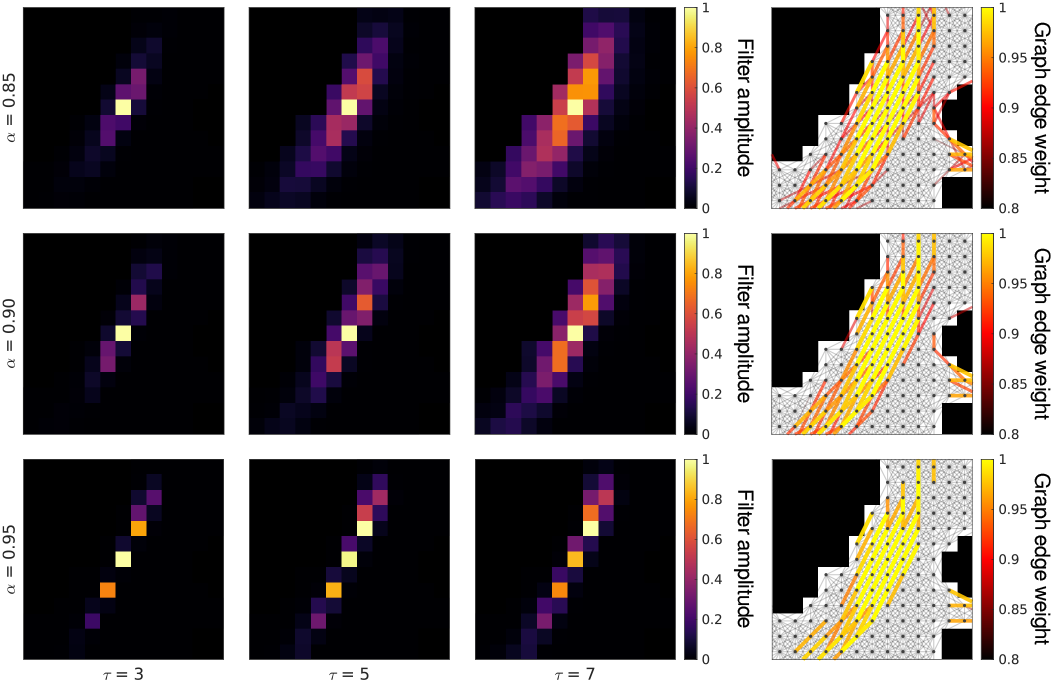
Effects of parameters τ and α on the shape of DSS filters located at red ROI shown in Figure 5. Graph parameters: 5-conn neighborhood, *β* = 50. Filters are shown normalized to the [0, 1] range.

The choice of neighborhood definition plays a significant role in the shape of the resulting filters. In combination with the 5-conn neighborhood definition, higher α values can result in non-local averaging filters when the ODFs are oriented along a neighborhood direction in the outer shell of the neighborhood (see Figure 5 middle left, Figure 6 bottom row). This effect is not present in filters created using the 3-conn neighborhood definition (see Figure S2), which additionally show a more limited capacity to represent orientation due to the reduced angular resolution of the neighborhood definition. More exhaustive results for both 5-conn and 3-conn filters are presented in Supplementary Figures S1-S6.

### 3.2. Validations on circular phantoms

Circular phantoms of 93 different orientations and 3 different radii were created as described in Section 2.5. Each phantom was corrupted with 10 realizations of additive white Gaussian noise of standard deviation 1, and subsequently denoised by spatial filtering with GSS and DSS over a range of parameters. The FWHM of GSS and the τ parameter of DSS were varied over a range from 1 to 8 in unit steps. Both the 3-conn and 5-conn neighborhood definitions were tested for DSS, which we will refer to as DSS3 and DSS5, respectively. The α parameter of DSS was set to 0.9 throughout.

To assess the denoising performance of GSS, DSS3 and DSS5, we performed receiver operating characteristic (ROC) analyses. The filtered phantom volumes were each thresholded at 300 uniformly-spaced consecutive levels spanning the minimum and maximum value in each filtered volume. The resulting detections for each threshold level were compared with the ground truth of the phantom, yielding true positive rates (TPR) and false positive rates (FPR) that were collected in ROC curves. The area under the curve (AUC) of the ROC curves was then computed, resulting in an overall measure of performance. Figures 7(a) and (b) show the overall performance of DSS3, DSS5 and GSS as characterized by the AUCs. Due to the lack of equivalence between DSS and GSS filters, there is no direct correspondence between individual values of FWHM and τ. However, it can be noted that the performance of GSS peaks at 2 mm FWHM, and diminishes for larger filter sizes. On the other hand, both DSS3 and DSS5 achieve substantially higher maximum performances, which are not negatively affected by increased filter size in the range of τ tested.

**Figure 7:**
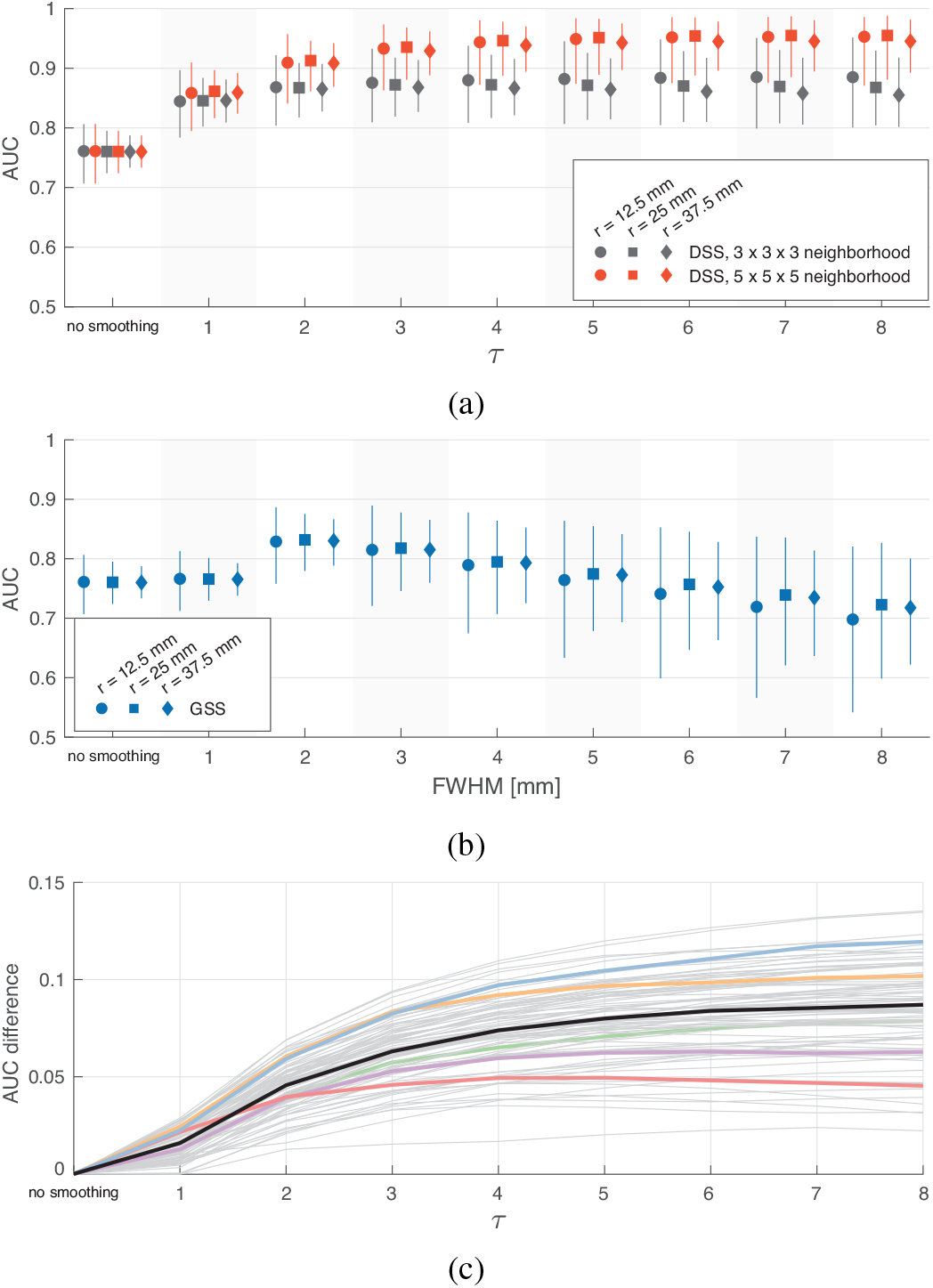
Validation of spatial smoothing on circular phantoms. (a)-(b) AUC of ROC curves obtained from volumes spatially smoothed with DSS and GSS, respectively. The markers show the median AUC over 930 ROCs (93 orientations × 10 realizations), whereas the whiskers represent 5 − 95% percentiles. (c) Difference between AUC values for DSS5 and DSS3 for phantoms with 25 mm radius. The black curve shows the difference between the median performances shown in (a), whereas the remaining curves show the difference between the 10-realization medians for each of the 93 phantom orientations. The five colored curves correspond to the phantom orientations shown in Fig. 4(a).

The median AUC of DSS5 consistently falls above that of DSS3 for τ ≥ 2 and all three phantom radii. The performance gap between DSS5 and DSS3 increases for larger τ, and slightly increases on circular phantoms with larger radii, i.e., smaller curvatures. These results corroborate the improvements in detection performance thanks to the increased angular resolution of the 5-conn neighborhood definition. This is further illustrated by Figure 7(c), which shows the performance improvement of DSS5 over DSS3 for individual phantoms orientations. The wide range of performance gains is representative of the varying difficulty of representing specific spatial orientations in the discrete domain of graphs, highlighting the importance of angular resolution for the proposed filters.

Given the overall superior performance of DSS5 over DSS3, in the following, DSS results are only presented for graphs using the 5-conn neighborhood definition.

### 3.3. Validations on streamline-based phantoms

A similar analysis was performed on streamline-based phantoms. A single phantom with *N*_*s*_ = 50, 100 and 200 streamline activations was created for each of the 95 subjects as described in Section 2.6. As in the analysis on circular phantoms, each phantom was corrupted with 10 realizations of additive white Gaussian noise of standard deviation 1, and denoised by spatial filtering with GSS and DSS over the same range of parameters. The α parameter of DSS was set to 0.9, whereas values of 0.85 and 0.95 were also tested on the 100-streamline phantoms. The denoising performance of both methods was assessed by applying the same ROC/AUC analysis described in Section 3.2.

Figures 8(a) and (b) show AUC results on all three types of phantoms for DSS and GSS, respectively. Due to the substantial amount of noise present in the phantoms, spatial smoothing using either GSS or DSS generally leads to better performance compared to no smoothing. DSS outperforms GSS across the range of τ and FWHM values tested, and across the different settings. As with the circular phantoms, the performance of GSS peaks at 2 mm FWHM, with increased size negatively affecting performance beyond that value. DSS shows a similar pattern, with peak performance achieved for τ of 3 and 4 for α = 0.9. Both GSS and DSS show better performance on phantoms with a greater number of streamlines. Additional results show that DSS outperforms GSS in both sensitivity and specificity (see Supplementary Figure S7(a)), and across a range of SNR values (see Supplementary Figure S7(b)).

**Figure 8:**
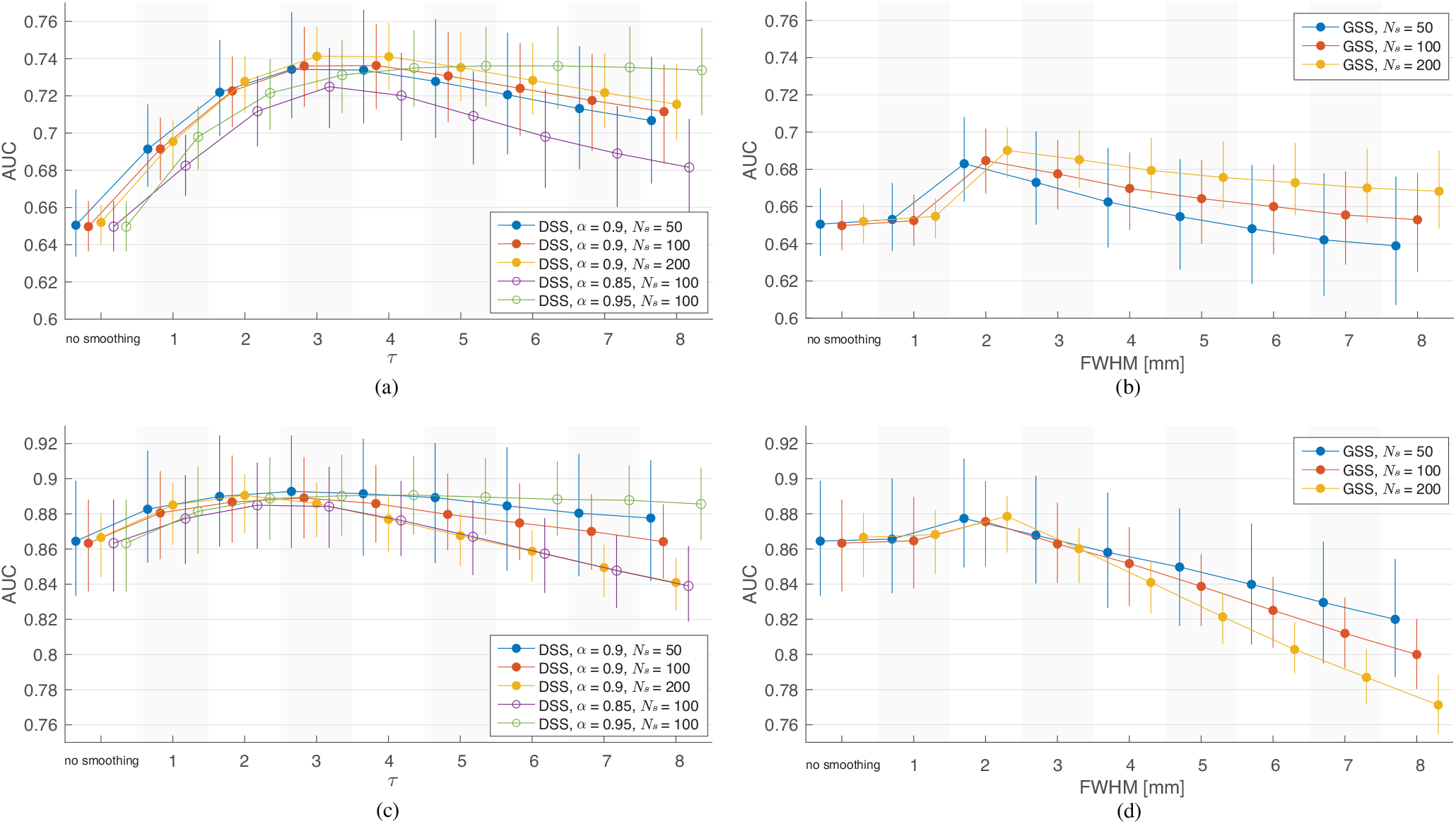
Validation of spatial smoothing on streamline-based phantoms. (a)-(b) AUC of ROC curves obtained from volumes spatially smoothed with DSS and GSS, respectively. (c)-(d) AUC of ROC curves obtained from activation mapping t-maps of time-series streamline-based phantoms smoothed with DSS and GSS, respectively. The markers show the median AUC over 950 ROCs (95 subjects × 10 realizations), whereas the whiskers represent 5 − 95% percentiles.

To assess the performance of DSS and GSS in combination with temporal modeling, i.e., as used within fMRI activation mapping studies, time-series version of the streamline-based phantoms were generated as described in Section 2.6. The phantoms were corrupted with additive white Gaussian noise of standard deviation 1 and subsequently spatially filtered with GSS and DSS with the same range of parameters used previously. The smoothed phantoms were subjected to a standard single-subject analysis in SPM, and the resulting t-maps were used in the ROC/AUC analysis.

Figures 8(c) and (d) show AUC results from the time-series phantoms. Due to the increased detection power afforded by temporal modeling, AUCs are higher for all scenarios in the time-series analysis compared to those in the single-volume analysis. Similarly to the single-volume phantom results, GSS achieves its best performance for 2 mm filters, and considerably deteriorates beyond that size. Notably, GSS only provides a distinct improvement over no smoothing for 2 mm filters. DSS results also show a negative correlation between filter size and performance for τ > 2, but the overall performance is superior to GSS and provides a benefit over no smoothing in most tested cases, with best results achieved for τ between 2 and 4. After subjecting the t-maps to activation mapping with false discov ery rate (FDR) correction at 5% (Genovese et al., 2002), the detection maps resulting from DSS showed substantially higher sensitivity and specificity than those from GSS (see Supplementary Figures S8-S10). These results also illustrate that the diminished performance of both methods on phantoms with a greater number of streamline activations is a consequence of increased FPR when using large filters.

Figures 8(a) and (c) also illustrate the effects of varying the α parameter of DSS in single-volume and time-series phantoms, respectively. For both types of phantoms higher values of α generally resulted in better performance. In the case of single-volume phantoms, filters with α=0.9 outperformed the others for small filter sizes, while α=0.95 is superior for larger filter sizes and across all sizes for time-series phantoms. In addition, filters with α = 0.95 show minimal decay in performance as filter size increases for both versions of the phantoms. Filters with α = 0.85 consistently performed worse than the others.

### 3.4. Single-subject task fMRI results

In order to explore the effects of the proposed smoothing method on real task fMRI data, we used SPM12 to perform activation mapping on the HCP100 task fMRI data, comprising 23 experimental conditions across 7 tasks. Each GLM analysis included 12 motion regressors (raw and temporal derivative) in addition to regressors for 2 to 8 experimental conditions associated with each task. The canonical HRF model, corresponding to a double gamma, was used—although such a temporal model is not tailored to the WM BOLD signal, it affects GSS and DSS equally, and should have no discernible influence on spatial filtering comparisons. Temporal noise modeling was done using a global AR(1) model. The fMRI data were smoothed using GSS and DSS with the same parameters used previously. For GSS, each fMRI volume was first multiplied with the WM mask, to avoid introducing signal from GM. This step is not required for DSS, as the method by its nature functions only in WM. The resulting t-maps were then thresholded to determine significant active voxels after FDR correction at 5%. Our choice of FDR as the correction method was due to it only assuming the *p*-values to be uniformly distributed under the null hypothesis. Correction methods based on assumptions about the smoothness of the data, such as those based on Gaussian random field theory, would be difficult to justify for an adaptive smoothing approach.

The sheer number of detection maps generated by this analysis—37,145 maps (95 subjects × 23 conditions × 17 filter settings)—renders exhaustive visual examination of them impracticable. Therefore, in our presentation, we focus on representative results that highlight the differences in maps generated by GSS and DSS. The full set of unthresholded t-maps is made available at NeuroVault^5^.

Figure 9 shows representative t-maps and detections from two subjects generated by DSS and GSS, with unmasked (i.e., full brain) GSS results included for reference.^6^ Visual inspection of the t-maps reveals that GSS results in generally round features with little oriented structure, with very little visible structure remaining for larger Gaussian filters. In contrast, t-maps obtained using DSS present notable spatial detail, with linear features in the shape of streamlines discernible across filter sizes. These differences are also present in the detection maps from both methods. While GSS detections are generally large and rounded—with very few detections present for smaller filters—DSS manifests detection maps with pronounced subtle spatial details—with considerable detections even for small filter sizes. The detections presented in Figure 9(a) highlight the capability of DSS in identifying separate streamline-shaped activations in two contiguous parallel axonal bundles (orange arrow), which remain distinct across the tested filter sizes. On the other hand, with GSS, these activations are combined into a single active region when large filters are used, and are not present when small filters are used. Notably, the case of FWHM = 3 mm shows activation foci being combined *across* rather than along axonal bundles, suggesting that these activations may not be separable with GSS. In Figure 9(b), DSS activation maps manifest an elongated, clearly resolved streamline-shaped activation that spans the corpus callosum (orange arrow), which is mostly undetected in GSS activation maps. In addition, the activations seen around the edges of the WM mask deserve notice. Although these activations may be attributed to interpolation artifacts or partial volume effect, due to them consistently being found in positions adjacent to active GM regions, it is important to note that both GSS and DSS produce these activations solely on the basis of signal from WM. DSS generally manifests more such activations, especially for small filter sizes. Additional activation mapping results are presented in Supplementary Figures S11 and S12.

**Figure 9:**
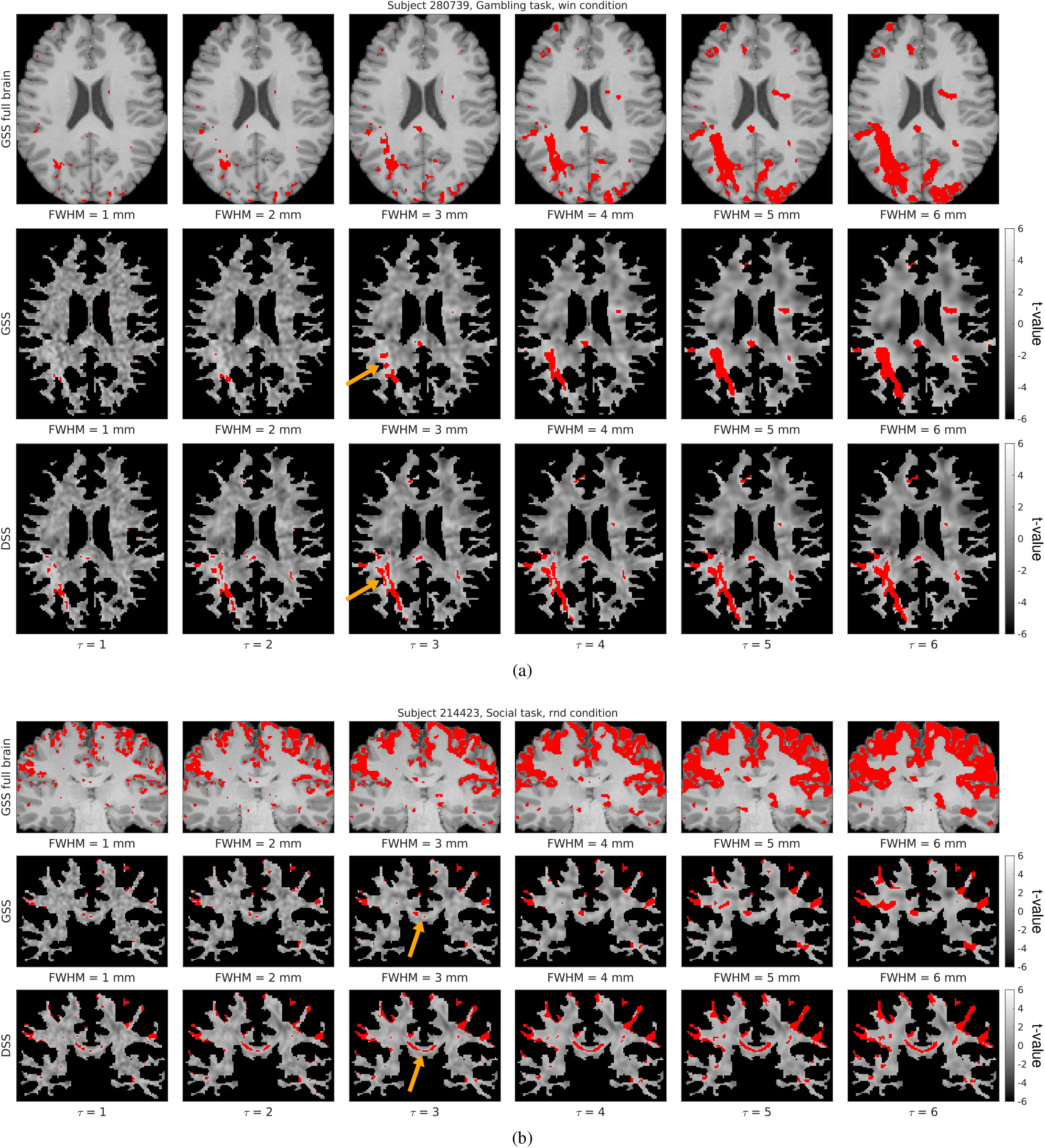
Comparison of representative single-subject activation mapping results generated with GSS and DSS, with t-maps shown in grayscale and detections overlaid in red (FDR-corrected at 5%). Full-brain activation maps are also shown for reference, overlaid on the subject’s T1w image.

In order to quantitatively investigate the degree to which spatial structure is present in t-maps obtained using the two smoothing methods, we analyzed the t-maps using *structure tensor* methods (Knutsson, 1989). While a thorough introduction to such methods falls outside the scope of this work, it is sufficient for our purposes to point out that the eigenvalues and eigenvectors of the structure tensor provide information on the presence and orientation of spatial structure, in the form of lines and edges, at a given point in an image or volume.

For each t-map, we constructed a quantitative structure map by computing the sum of the structure tensor eigenvalues at every voxel (a measure of the amount of spatial structure in each voxel). The mean value of each structure map provides a global measure of the presence of spatial structure in the corresponding t-map. Figure 10(a) shows a comparison of this global structure measure for DSS and GSS. For both methods the amount of structure present in the t-maps diminishes as the filter size increases, which is consistent with the loss of spatial detail resulting from smoothing the data. This effect is very pronounced for GSS, while t-maps generated using DSS exhibit a more consistent amount of spatial structure across the tested filter sizes.

**Figure 10:**
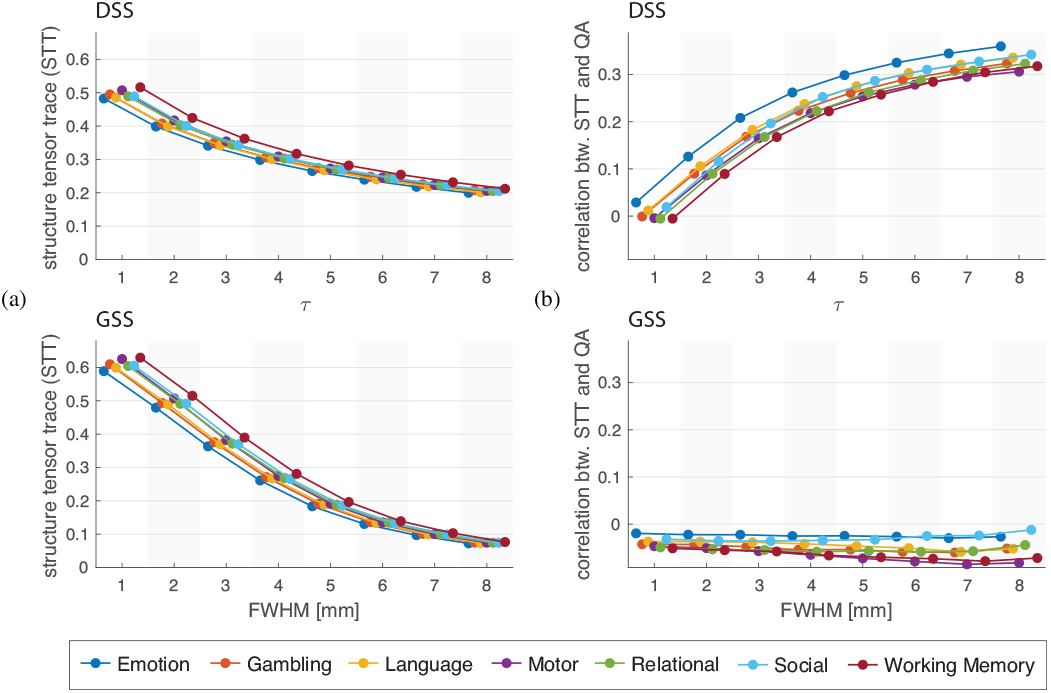
Structural analysis of task fMRI t-maps, obtained using local structure tensor analysis (Knutsson, 1989) where the eigenvalues of the structure tensor denote the amount of spatial structure. (a) Quantification of the amount of anisotropic structure observed in t-maps, specified by the mean structure map value, averaged across the task’s experimental conditions. (b) Correlation between subjects’ QA maps and structure maps, averaged across each task’s experimental conditions. Markers shows the median value across the 95 subjects.

To determine the extent to which the structure present in the t-maps is influenced by the diffusion information introduced by DSS, we computed Pearson’s correlation coefficient between the quantitative structure maps and the quantitative anisotropy (QA) map (Yeh et al., 2013) of the associated subject; see Figure 10(b). For DSS, this correlation is close to zero at τ = 1, and steadily increases for increased filter sizes. In contrast, the structure manifested in t-maps obtained through GSS shows a slightly negative correlation with QA, which stays nearly constant across all filter sizes. These results suggest that DSS is successful at informing the smoothing process with the local diffusion properties of the underlying WM, with larger values of τ resulting in stronger diffusion encoding.

Figure 11 compares the number of detections obtained from DSS and GSS. To prevent bias due to differences in brain size, we present the fraction of each subject’s WM mask being declared as active. Overall, the detection rates for both methods increase as a function of filter size, with DSS exhibiting a more linear increase than GSS. While the number of detections on t-maps obtained from volumes smoothed with DSS and GSS is comparable for large filters, DSS generally produces substantially more detections with smaller filter sizes, as manifested by comparing the median detection numbers of corresponding tasks.

**Figure 11:**
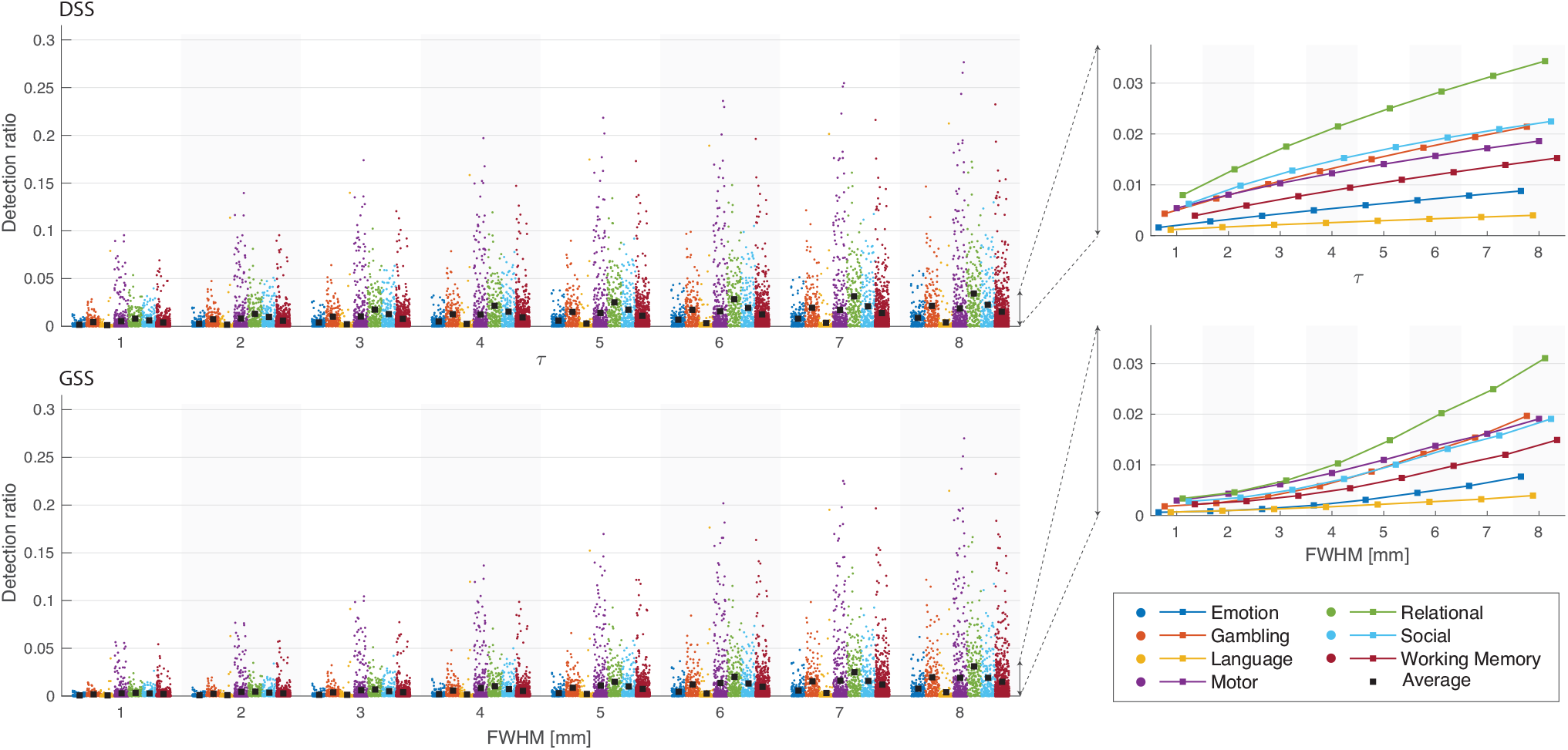
Fraction of voxels within WM mask detected as being significant using DSS (top left) and GSS (bottom left) across 7 functional tasks, over 95 subjects. Significant voxels were determined after FDR correction at 5%. In the plots on the left, each dot corresponds to one subject, whereas ▪ shows the median value across the 95 subjects. The plots on the right show the trend of the average value as a function of filter parameters τ and FWHM for GSS and DSS respectively.

In the absence of ground truth, it is not possible to make definitive statements on the relationship between differences in the number of voxels deemed active by each method and potential differences in their sensitivity and specificity. However, it can be insightful to quantify the difference between the detection maps generated with DSS and GSS. To quantify the similarity between a pair of detection maps we computed the Dice coefficient between them, defined as

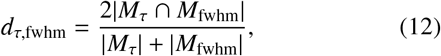

where *M*_τ_ denotes the set of detected voxels using DSS with a given τ, *M*_fwhm_ denotes the set of detected voxels using GSS with a given FWHM, and | · | denotes set cardinality. The Dice coefficient is constrained to the [0, 1] range, where a value of 1 signifies perfect overlap between the detection maps and a value of 0 represents no overlap.

For every subject and experimental condition we calculated Dice coefficients between detection maps obtained with GSS and DSS of all filter sizes, and arranged them into 8× 8 Dice matrices. Additionally, we calculated the maximum Dice coefficient between each DSS filter size and every GSS filter size for each subject and condition. Figure 12 shows Dice results for several representative experimental conditions. The overall similarity between the detection maps obtained with DSS and GSS is relatively low. The highest ensemble Dice is achieved for τ = 7 and FWHM = 8 mm, where it reaches a value of 0.65, with other combinations achieving values close to this one (see ensemble Dice matrix). The relationship between the τ and FWHM values that result in the highest similarity in the detection maps is also shown to be nonlinear, tracing a particular curve across the Dice matrices that is generally similar across experimental conditions. The similarity between the detection maps also shows considerable variation across tasks and individual experimental conditions (see results for all experimental conditions in Supplementary Figure S13), with below-average similarity in the Language and Motor tasks and above-average in the Gambling and Relational tasks.

**Figure 12:**
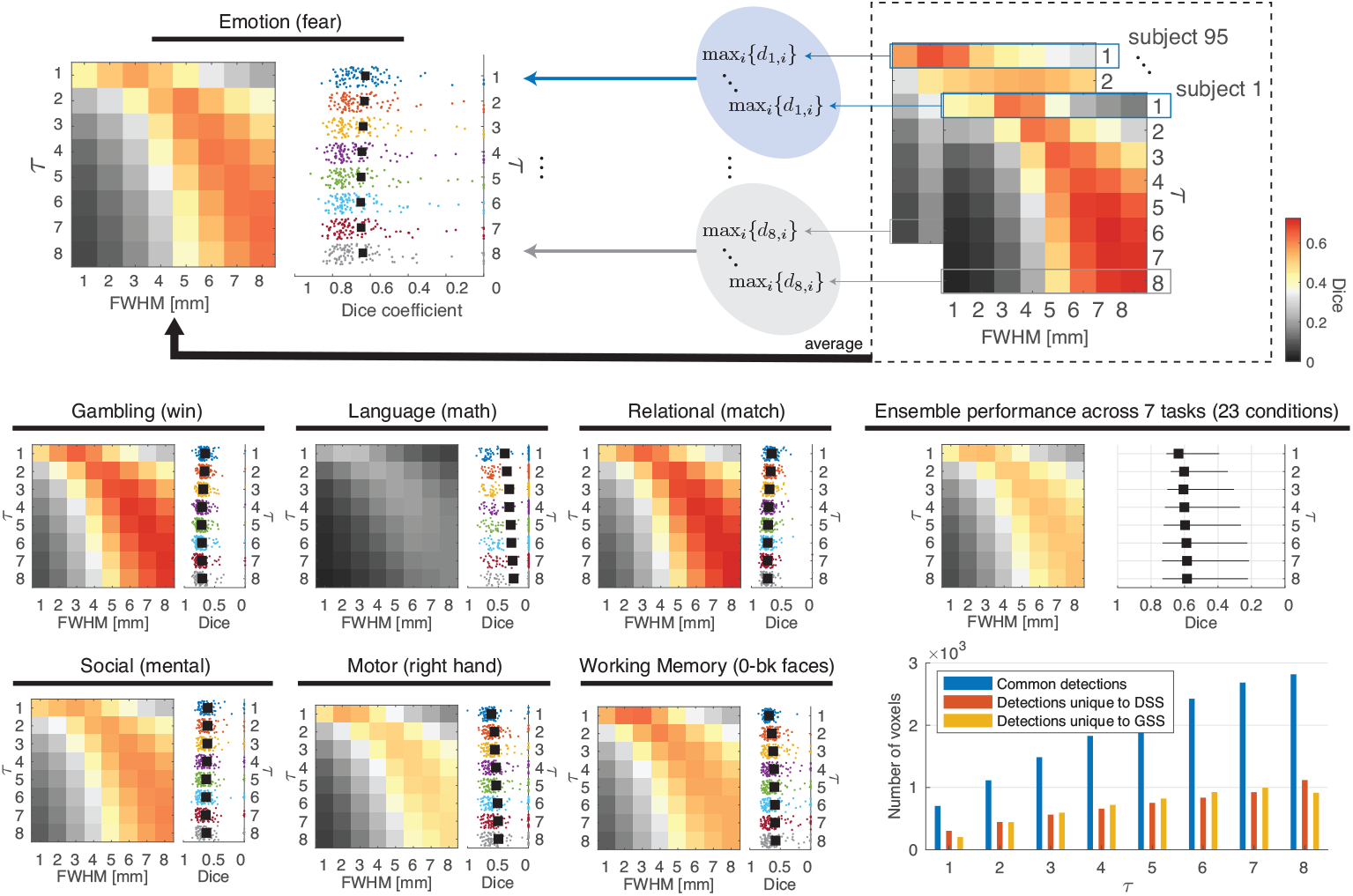
Dice similarity between detection maps generated with DSS and GSS. For each subject and condition, an 8 × 8 Dice matrix was computed, where each element represented *d*_τ,fwhm_, see (12). For a given subject, if neither DSS nor GSS led to any detections for a given combination of τ and FWHM, the corresponding element was excluded from further analysis. The schematic on top explains how the results were ensembled across subjects, resulting in two plots for each experimental condition; in the plots on the right, the mean of the scattered values is indicated by ▪. Results are presented for a representative experimental condition in each task—see results across the 23 conditions in Supplementary Figure S13, as well as ensembled across 23 conditions; the ensemble plot on the left shows the average across conditions, whereas the one on the right shows the median and range of the mean maximum Dice values across conditions. The plot on the bottom right shows the average number of common and unique detections generated by DSS and GSS across all subjects and conditions, wherein every value of τ was compared with the FWHM that resulted in the maximum Dice coefficient.

In order to determine whether the detections generated by either method are a subset of the detections from the other, we examined the number of common and unique detections produced by DSS and GSS. For all subjects and experimental conditions, the detection maps produced by DSS were compared with the most similar maps produced by GSS. Figure 12, bottom right, shows the average number of voxel detections common to both methods, as well as those unique to each method, for the tested values of τ. These results show that, across filter sizes, both DSS and GSS produce a considerable number of detections that are not produced by the other method. This observation, together with the generally low Dice similarities, suggests the presence of substantial differences in the localization and spatial extent of activations detected using DSS and GSS.

### 3.5. Group task fMRI results

We performed random-effects group analysis based on the single-subject results for each of the 23 experimental conditions across the seven tasks. The estimated regressor weights of each experimental condition were taken to MNI space using the displacement maps provided with the HCP data—the inverse of those used to map the preprocessed fMRI data to ACPC space—and a GLM was fitted to them to create group t-maps. These group maps were then thresholded to determine significant active voxels after Bonferroni correction at 5%.

Figure 13 shows representative results for one condition of the Gambling task. Overall, spatial patterns in the t-maps are more clearly visible than in the single-subject analysis, remaining more defined in the DSS results than in those of GSS. Interestingly, both methods show large WM regions in the shape of axonal bundles that are strongly anticorrelated with the experimental conditions.

**Figure 13:**
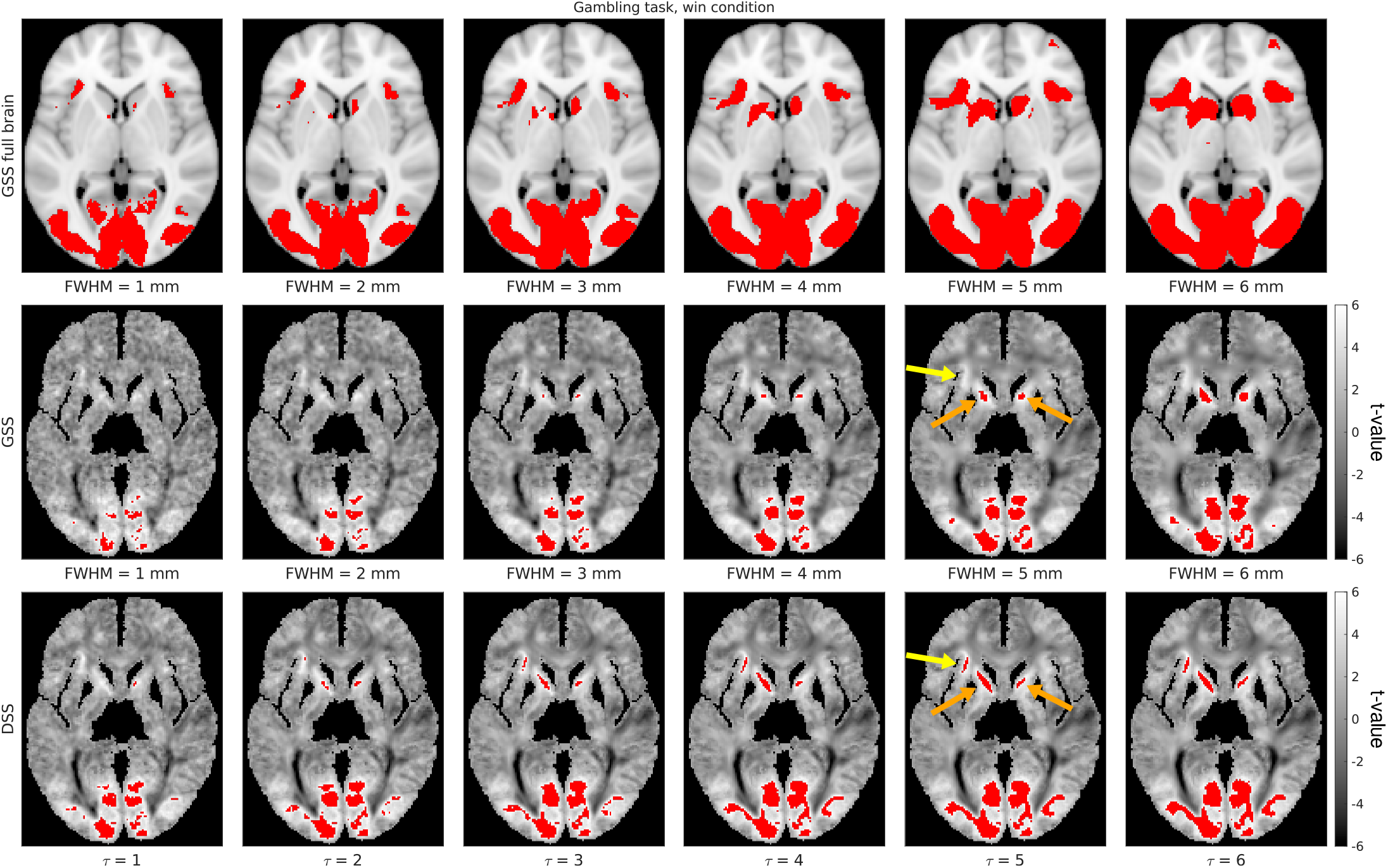
Comparison of representative group activation mapping results generated with GSS and DSS, with t-maps shown in grayscale and detections overlaid in red (Bonferroni-corrected at 5%). Full-brain activation maps are also shown for reference, overlaid on the MNI152 T1w template image.

The activation maps in Figure 13 show similar patterns to the single-subject activation maps. While DSS is capable of producing elongated, streamline-like detections, those of GSS are generally round. In addition, DSS reveals considerable detections for small filter sizes. Additional group activation mapping results are shown in Supplementary Figures S14-S16.

In order to study the consistency of the results obtained by each method, we investigated the test-retest reliability of GSS and DSS through a Monte Carlo experiment. The 95 subjects were repeatedly split into two groups, after which a random-effects model was fitted to each group, and the resulting t-maps and detection maps were compared. This process was repeated 30 times, and the similarities of the resulting t-maps and detection maps were quantified using Pearson correlation and Dice similarity, respectively. Figure 14 shows results of this analysis for a representative subset of experimental conditions. Correlation and Dice scores show an increasing trend with respect to the filter size, for both GSS and DSS. The values produced by both methods are roughly comparable, being slightly higher overall for DSS, particularly for small filter sizes. Full comparisons for all experimental conditions are presented in Supplemetary Figure S17.

**Figure 14:**
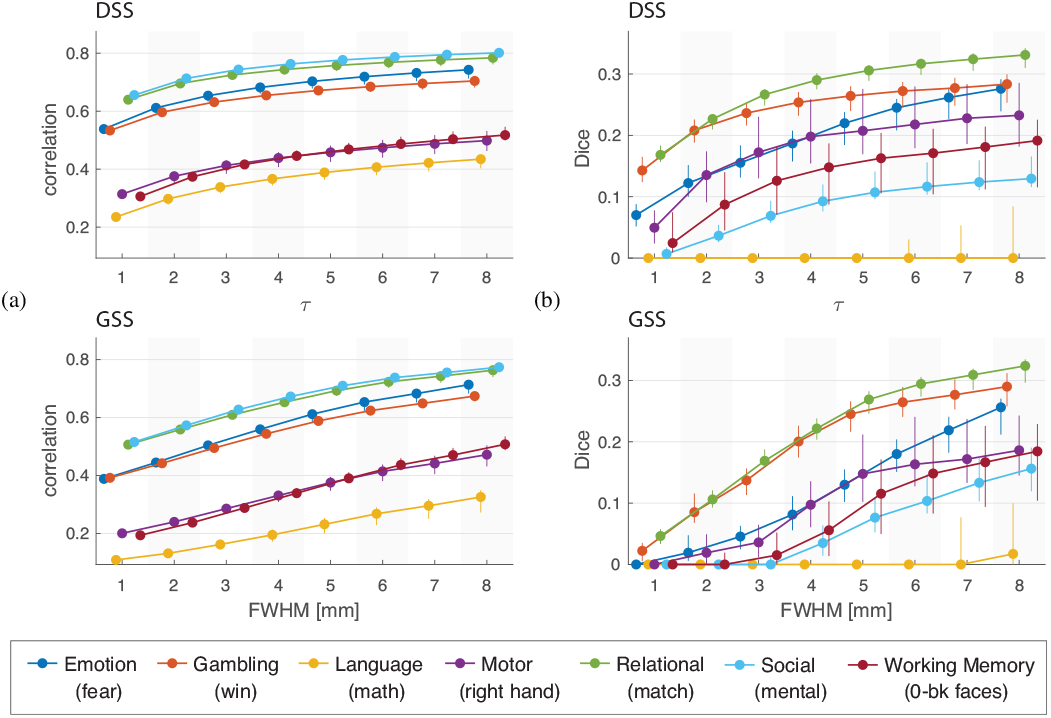
Results of Monte Carlo test-retest analysis for one representative experimental condition from each task. Subjects were repeatedly divided into two groups and subjected to group analysis, and the resulting statistical maps were compared. (a) Correlation between t-maps of both groups. (b) Dice similarity between activation maps of both groups. The markers show the median value across 30 experiments, whereas the whiskers represent 5 − 95% percentiles.

### 3.6. Processing time

Although the proposed methodology requires additional MRI scanning time for the acquisition of DW-MRI data, it does not impose a dramatic increase in processing time over conventional approaches. Using a workstation with an Intel Core i7-7700K processor and 64 GB of RAM, the generation of diffusion ODFs from DW-MRI data required approximately 90 seconds. The graph and its Laplacian matrix could then be calculated from the ODFs in under 15 seconds. Both of these operations need only be performed once per subject.

In our implementation, the average filtering time of a single volume with GSS was 10.3 ms using the imgaussfilt3 MATLAB function (the same operation required about 450 ms when using the smoothing implemented in SPM). On the other hand, DSS filtering scales efficiently with the number of filter kernels used. Average single-volume DSS filtering times for a single kernel were 115 ms for the 5-conn neighborhood and 56 ms for 3-conn, and became reduced to 17.7 ms and 11.0 ms, respectively, when using 8 filter kernels at once. With worst case performance, the proposed method gave filtering times of around 45 seconds for a 405-volume series (the longest of those available in HCP data, corresponding to the Working Memory task).

## 4. Discussion

### 4.1. Interpretation of results from simulated data

Previous implementations of voxel-wise graphs on GM (Behjat et al., 2015; Maghsadhagh et al., 2019; Behjat and Larsson, 2020) have used the 3-conn neighborhood in defining graph edges. However, given the different nature of the proposed encoding for WM graphs—representing axonal orientations rather than GM morphology, we considered the potential advantages of using a larger neighborhood definition. To this end, we compared the denoising performance obtained with graphs using the 3-conn and 5-conn neighborhood definitions on circular phantoms of multiple orientations and radii. Such phantoms were used because, barring discretization artifacts, they offer an exhaustive sampling of all possible orientations in which data can appear in three dimensions. The results show a clear improvement from using the larger neighborhood definition (see Figures 7(a) and (c)), which can be attributed to its superior angular resolution of 98 neighborhood directions, against the 26 of the 3-conn definition. Furthermore, comparing performances obtained on phantoms of different radii shows that the larger neighborhood definition provides more stable performance across spatial curvatures than the smaller neighborhood, which performs worse for smaller curvatures, particularly for larger filters. Compared to isotropic Gaussian smoothing (see Figure 7(b)), both the 3-conn and 5-conn neighborhood definitions used in DSS showed enhanced denoising performance on circular phantoms. In particular, while the performance of GSS deteriorates for larger filter sizes, the performance of DSS reaches a plateau instead, suggesting that the diffusion-informed nature of DSS filters is capable of minimizing the introduction of spurious signal even for larger filter sizes.

To better mimic spatial activation patterns manifested as BOLD contrast in WM, we designed and studied semi-synthetic streamline-based phantoms, whose diffuse activation patterns are representative of WM fiber structures, along which correlated BOLD activity is expected (Ding et al., 2013, 2016). The phantoms were studied in two settings. In the first setting, the denoising performance was studied in the absence of temporal modeling, wherein both methods provided an improvement over no smoothing, but DSS outperformed GSS for all tested filter sizes (Figure 8(a)). In the second setting, the phantoms were studied within the context of GLM activation mapping, i.e. with temporal modeling, wherein GSS provided only minimal improvements over no smoothing, whereas DSS provided a notable improvement (Figure 8(b)). In addition, when the timeseries phantoms were subjected to activation mapping with FDR correction, activation maps from GSS showed reduced sensitivity and specificity when compared to those of DSS (see Supplementary Figures S8-S10). The phantoms were also used to study the influence of the α parameter of DSS, which sets a lower bound on the weight of connections allowed in the WM graph. Due to the narrower and more directional filters resulting from higher α values (Figure 5, Supplementary Figures S1-S6), the increased performance on the streamline-based phantoms would be expected (Figures 8(a) and (c)). However, this result may not be readily extensible to real fMRI data, as the spread of real activation patterns is not known.

### 4.2. Interpretation of results from real data

We compared single-subject activation mapping results from DSS and GSS on task fMRI data from the HCP100 subject set. Structure tensor analysis of the resulting t-maps revealed that the overall amount of structure present diminished for larger filter sizes, an effect that is more pronounced for GSS (Figure 10(a)). Such results reflect the loss of spatial details that happens as a result of lowpass filtering. However, due to the highly anisotropic shapes that DSS filters take within the WM (Figure 5), features in the shape of lines and edges can be present in t-maps even for larger filter sizes (Figure 9). In addition, the spatial structure present in the t-maps obtained with DSS is correlated with regions of high diffusion anisotropy (Figure 10(b)), indicating that DSS successfully adapts its smoothing to the underlying WM microstructure.

Due to the differences in their definitions, as well as the adaptive nature of DSS, there is no direct correspondence between GSS and DSS filters. This is corroborated by the relatively low Dice coefficients between detection maps resulting from both methods (see Dice matrices in Figure 12). The overall number of detections is comparable for GSS and DSS, with a considerable and roughly equal number of activations being unique to each method (see bar chart in Figure 12, bottom right). On the other hand, example detection maps corroborate that DSS is capable of resolving subtle, slender activation patterns along axonal pathways across multiple filter sizes by leveraging information about the spatial correlation structure of the BOLD signal in WM. Figure 9(a) exemplifies the increased resolution from DSS, presenting a case where it is capable of resolving two parallel streamline-like activations that GSS is incapable of identifying as separate. Figure 9(b) illustrates a similar case, with DSS detecting a highly resolved streamline-like activation through the corpus callosum that is left largely undetected by GSS. Supplementary Figures S11 and S12 present additional detection map comparisons highlighting the increased sensitivity and specificity of the proposed methodology over conventional GSS.

We also compared group activation mapping results from DSS and GSS. Similarly to the single-subject results, group t-maps obtained with DSS manifested intricate spatial structures across filter sizes, while t-maps generated with GSS presented mostly smooth, round features (Figure 13). The same patterns extended to the activation maps produced by both methods, where DSS has shown greater specificity and an increased number of detections in multiple instances (Supplementary Figures S14-S16). Although group WM activations obtained with GSS and DSS are often contained within those obtained with full brain GSS, it is important to note that while the latter rely mostly on signal from the GM, the former rely solely on signal from the WM, and result in much greater specificity in the detected activations.

In order to evaluate the consistency of the statistical maps generated by both methods, we performed a test-retest analysis of group activation mapping. While the performances of DSS and GSS were comparable for the upper range of filter sizes tested, DSS showed a marked improvement for small filter sizes (Figure 14), altogether suggesting that DSS is capable of yielding equally or more consistent results than GSS is.

### 4.3. Limitations

We used a sigmoid function, see (10), as a means of boosting orientation encoding, allowing diffusion only along main directions of diffusion coherence. We studied three threshold values, α = 0.85, 0.9 and 0.95, all of which yielded better performance than GSS on phantom data, with noticeable variations in performance among the three values. However, the general choice of the thresholding function and its associated parameters is rather ad-hoc, which is a complication of similar nature as that encountered in connectomic studies (Rubinov and Sporns, 2010). Future work should consider a more rigorous validation of the thresholding scheme for obtaining optimal performance, especially on real fMRI data.

Accurate co-registration of functional, structural, and diffusion MRI data is a cornerstone of the proposed methodology. Within this study, we used preprocessed HCP data, which have been diligently motion-corrected, distortion-corrected, and coregistered (Glasser et al., 2013). However, conducting solid preprocessing steps may not be possible in some datasets, and if so, results obtained using the proposed method on such datasets should be interpreted with care.

A number of recent studies have highlighted substantial differences between the HRF in WM and that in GM (Li et al., 2019b; Wang et al., 2020b; Choi et al., 2020), which corroborate similar sporadic observations from earlier studies that showed evidence for delayed and subdued hemodynamic responses compared to that in GM (Yarkoni et al., 2009; Fraser et al., 2012), and in particular, in the corpus callosum (Tae et al., 2014; Courtemanche et al., 2018). The recent evidence for the unique features of HRF in WM is indeed insightful, but given the ongoing nature of this research, we decided to use the standard HRF model that is conventionally used in fMRI activation mapping in the present work. Given that our work is comparative, the choice of the HRF model affects both DSS and GSS equally, and as such, we do not believe that our conclusions would be substantially affected by the use of a more precise model. Nevertheless, future work aimed at investigating the BOLD signal in WM can most likely benefit from combining a more appropriate HRF model with adaptive smoothing of the BOLD signal by DSS.

### 4.4. Outlook; potential extensions and other applications

Due to the limited degree to which diffusion ODFs can differentiate fiber orientation (Jones et al., 2013), we boost orientation encoding by means of a weight thresholding scheme. Alternatively, the proposed design can be extended to leverage standard fiber orientation distribution (FOD) functions estimated from either the diffusion ODFs (Descoteaux et al., 2008) or the raw diffusion data (Tournier et al., 2007), or asymmetric FODs (Bastiani et al., 2017), to obviate the need for thresholding. In the absence of HARDI data but presence of DTI data, the proposed method can be readily extended to leverage diffusion tensors instead of diffusion ODFs, e.g. as in Tarun et al. (2019), which can be of particular interest for reanalyzing the vast extent of currently available fMRI datasets that are accompanied by DTI data.

In the absence of any DW-MRI data, it would be possible to adapt the proposed method to use a structure tensor representation (Knutsson, 1989) derived from T1-weighted MRI images as the complementary contrast (Abramian et al., 2020b), wherein the proposed filtering scheme could be extended to function across the entire brain mask. The resulting morphology-based spatial smoothing could then be seen as a GSP-based alternative to non-linear filtering algorithms which enable spatial smoothing within similar anatomical compartments (Smith and Brady, 1997; Weickert and Scharr, 2002; Ding et al., 2005; Rydell et al., 2008; Lohmann et al., 2018), but will not provide adaptation to WM fiber orientations.

In addition to performing denoising through heat kernel smoothing (i.e., lowpass filtering), the proposed WM graphs can be used to implement graph-wavelet denoising, similar to that implemented by Behjat et al. (2015) for GM graphs, using novel data-driven GSP denoising schemes (de Loynes et al., 2019) in combination with computationally efficient multi-scale spectral graph decomposition methods (Li et al., 2019c; Shuman, 2020) that can be tractably implemented on large graphs.

In the present study, we only explored spatial smoothing of task-based fMRI data within the context of activation mapping, whereas DSS can be readily applied to WM resting-state fMRI data, where recent studies have used Gaussian smoothing of the data as a pre-processing step. Such research appears particularly promising in light of studies reporting the existence of BOLD-like response in resting-state data (Liu and Duyn, 2013; Petridou et al., 2013; Karahanoğlu and Van De Ville, 2015; Li et al., 2021), and the current growing interest in exploring functional dynamics of WM at rest (Peer et al., 2017; Ding et al., 2018; Li et al., 2019a; Wang et al., 2020a; Li et al., 2020a).

It is worth noting that DSS may prove beneficial for enhancing the detection of functional pathways through the use of functional-correlational tensors (FCT) (Ding et al., 2013) or high angular resolution functional imaging (HARFI) (Schilling et al., 2019). FCT and HARFI provide the means to derive functional WM pathways by characterizing the spatial anisotropy observed in the temporal correlation in the BOLD signal at adjacent WM voxels. Given the lack of spatial adaptiveness of GSS, its use is likely to distort the spatial anisotropy in the signal, on which these methods rely. On the other hand, filtering the fMRI data with DSS may help boost this spatial anisotropy, thus enhancing the detection of spatiotemporal correlation in the local BOLD signal. Furthermore, FCTs have been leveraged for improving inter-subject registration of resting-state data based on functional features (Zhou et al., 2018), which might also be enhanced if the data are initially filtered with DSS.

DSS may also be used as a method to filter tractography streamlines in a manner similar to SIFT (Smith et al., 2013). In particular, by applying DSS to voxelized representations of streamlines, the resulting filtered maps can be quantified to obtain a validity score for tracts—tracts that are closely aligned with the underlying diffusion map should be minimally deteriorated by DSS.

Another research avenue that can benefit from the proposed WM graph design is structural studies. The eigenvalues of cortical surface graphs as well as their eigenmodes have been leveraged in multiple applications, namely, quantifying cortical folding patterns (Germanaud et al., 2012; Rabiei et al., 2016; Dubois et al., 2019), age prediction (Wachinger et al., 2015; Masoumi et al., 2019), and analysis of brain asymmetry in health (Wachinger et al., 2015; Maghsadhagh et al., 2019) and in disease (Wachinger et al., 2016a,b; Masoumi et al., 2019). Such analyses can be extended to leverage the spectra of WM graphs. Analysis on similarly designed graphs using DW-MRI data—covering the entire brain rather than just the WM—has shown that an initial subset of the graph eigenmodes provides informative features to distinguish between subjects (Tarun et al., 2019). Lastly, ODF-based WM graphs may be found beneficial in deriving structural connectivity measures that account for direct as well as indirect pathways, for example, similar in nature to those derived from a recently proposed DTI-based conductance model (Frau-Pascual et al., 2019, 2020).

## 5. Conclusion

The development of methods geared specifically towards WM can prove substantially helpful in investigating the functional significance of the BOLD signal in WM. Notwithstanding the repository of sophisticated smoothing techniques found in the literature, to date, studies on fMRI data in WM have mainly resorted to isotropic Gaussian smoothing. An apparent reason is the ease in implementing Gaussian smoothing and its availability in widely used open-access software packages, which facilitate its routine application. The proposed diffusion-informed spatial filtering method, in conjunction with the use of WM-specific HRF models and MR sequences, holds promise to aid better understanding of the functional role of WM.

## Code and data availability

An implementation of the methods proposed in this work will be made available as a MATLAB package on GitHub. The simulated circular and streamline-based phantoms used in this work will be made available on the OpenNeuro platform. Single-subject and group activation mapping t-maps will be made available on the NeuroVault platform. Customized links will be added in the final version of the manuscript.

## Supporting information

Supplementary Results

## CRediT authorship contribution statement

**David Abramian:** Conceptualization, Methodology, Software, Data curation, Formal analysis, Writing - original draft, Writing - review & editing. **Martin Larsson:** Conceptualization, Methodology, Software, Writing - review & editing. **Anders Eklund:** Writing - review & editing, Supervision. **Iman Aganj:** Writing - review & editing, Supervision. **Carl-Fredrik Westin:** Writing - review & editing, Supervision. **Hamid Behjat:** Conceptualization, Methodology, Software, Validation, Writing - original draft, Writing - review & editing.

## Acknowledgements

Data used in this work were provided by the Human Connectome Project, WU-Minn Consortium (Principal Investigators: David Van Essenand Kamil Ugurbil; 1U54MH091657) funded by the 16 Institutes and Centers of the National Institutes of Health (NIH) that support the NIH Blueprint for Neuroscience Research; and by the McDonnell Center for Systems Neuroscience at Washington University.

This work, and HB, were supported by the Swedish Research Council under Grant 2018-06689, and in part by thán Foundation. DA and AE were supported by the Swedish Research Council under Grant 2017-04889, by the ITEA3 / VINNOVA funded project Intelligence based iMprovement of Personalized treatment And Clinical workflow supporT (IMPACT), and by the Center for Industrial Information Technology (CENIIT) at Linköping University. IA was supported by the BrightFocus Foundation (A2016172S) and the NIH, specifically the National Institute of Diabetes and Digestive and Kidney Diseases (K01DK101631) and the National Institute on Aging (R56AG068261).

A preliminary version of this work has been presented (Abramian et al., 2020a).

## Appendix A Frequency interpretation of graph Laplacian eigenvalues

In classical signal processing, in particular in the case of 1D discrete temporal signals, a set of complex exponentials *e* ^*jωx*^ of varying frequencies *ω* defines a basis that can be used to transform a given signal to a Fourier (spectral) representation. Importantly, these complex exponentials are the eigenfunctions of the one-dimensional Laplacian operator, i.e.,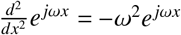. Given that a graph structure can be interpreted as a generalization of the 1D regular grid, the eigenvalues λ_*l*_ and eigenvectors **u**_*l*_ of the graph Laplacian **L** can be seen as analogous to the frequencies and complex exponentials of classical signal processing, respectively. With this interpretation, given two eigenvalues of **L** such that λ_*n*_ < λ_*m*_, it can be stated that the eigenvector associated with λ_*m*_ entails a notion of higher frequency—i.e., higher spatial variability—than the eigenvector associated to λ_*n*_. In the following we will illustrate this point in two ways.

Given a graph signal **f** ∈ 𝓁^2^(𝒢), the extent of variation of **f** on 𝒢 can be quantified by introducing a measure denoted as graph signal variation (GSV), defined as

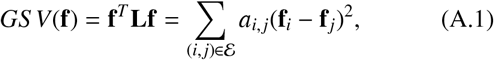

where larger values of *GSV*(**f**) represent greater variability of **f** on 𝒢. The eigenvectors of **L** can be equivalently seen as graph signals, and thus be quantified in relation to their extent of variation on 𝒢. By noting that i) the eigenvectors of **L** are orthonormal, i.e., 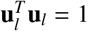 and ii) **Lu**_𝓁_ = λ _𝓁_ **u**_𝓁_, it follows that

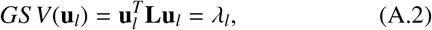

showing that the eigenvalue λ_*l*_ associated to each eigenvector **u**_*l*_ is a quantification of the extent of variability of **u**_*l*_.

The variability of eigenvectors can also be measured by examining their *zero crossings*—i.e., changes in their sign at adjacent graph vertices—using a weighted zero crossing measure (WZC) defined as

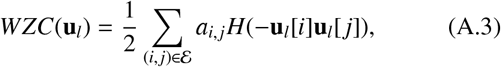

where *H*(·) denotes the Heaviside step function. To show the link between *WZC*(**u**_*l*_) and λ_*l*_, we calculated the WZC of an even sampling of 41 eigenvectors of **L** for fifty subjects— computing the full eigendecomposition of **L** is impractical due to its size. Figure A.1 shows the relation between λ_*l*_ and the *WZC*(**u**_*l*_), illustrating that larger eigenvalues entail a greater extent of spatial variability in their associated eigenvectors, as measured by the WZC. It should be noted that the monotonically increasing behavior of *WZC*(**u**_*l*_) relative to λ_*l*_, which holds up to the very upper parts of the spectrum, stops at the higher end eigenvalues. This is a consequence the decrease in delocalization manifested by eigenvectors of **L** at the upper part of the spectrum—unlike the complex exponentials of classical signal processing, which are delocalized, eigenvectors of **L** can present localized patterns of spatial variability. Nevertheless, given the lowpass profile of the spectral kernels used in this work, the loss of delocalization associated to the upper end of the spectrum is of no concern for the application at hand. For a more comprehensive overview of the link between classical signal processing and GSP, the interested reader is referred to Ortega et al. (2018); Stankovic et al. (2019); Huang et al. (2018a).

**Figure A.1:**
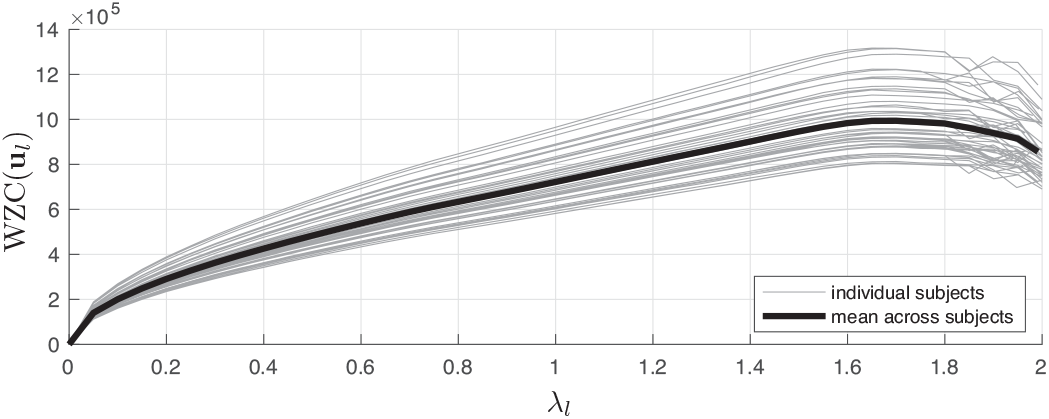
WZC of a subset of eigenvectors of the WM graph Laplacian of 50 subjects.

## Appendix B Spectral graph filtering through polynomial approximation

Spectral graph filtering can be efficiently implemented using polynomial approximation schemes (Hammond et al., 2011; Shuman, 2020), mitigating the need to diagonalize large **L** matrices as those used in the present work. Using this approach, a spectral kernel *k*(λ) is first approximated using a polynomial of suitable order, denoted *p*(λ) : [0, 2] → ℝ, and filtering of signal **f** is then implemented as

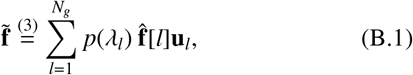

where the vectorized form of (3) is invoked. Noting that **Lu**_*l*_ = λ_*l*_**u**_*l*_ ⇒ *p*(**L**)**u**_*l*_ = *p*(λ_*l*_)**u**_*l*_, (B.1) can be simplified as

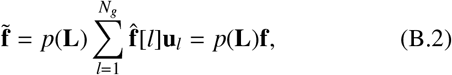

where in the last equality we used 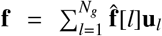. Using this scheme, filtering is performed through a series of polynomial matrix operations on **L**, without the need to access the Laplacian eigenvalues. In this work, we leveraged truncated Chebyshev polynomial approximations of spectral kernels as presented by Hammond et al. (2011), which have the benefit of approximating a minimax polynomial, minimizing an upper bound on the approximation error.

## Appendix C Uniform sampling of ODFs

We defined a spherical sampling grid using the vertices of an icosahedron with five levels of subdivision, which resulted in a total of 10,242 vertices on the unit sphere. Due to non-uniformity in the spatial spread of the vertices, the number and distribution of vertices that fall within the solid angles Ω_*i, j*_ subtended along the 26/98 different 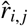 neighborhood directions vary. To overcome this bias, we treated the vertices that fall within Ω_*i, j*_ around the z-axis as a sampling template, resulting in *N*_*t*_ = 389 and 105 template directions for the 3-conn and 5-conn neighborhood definitions, respectively. The sampling template was then rotated and centered around each neighborhood direction 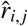, resulting in a set of sampling directions 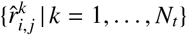 (see Figure C.1).

**Figure C.1:**
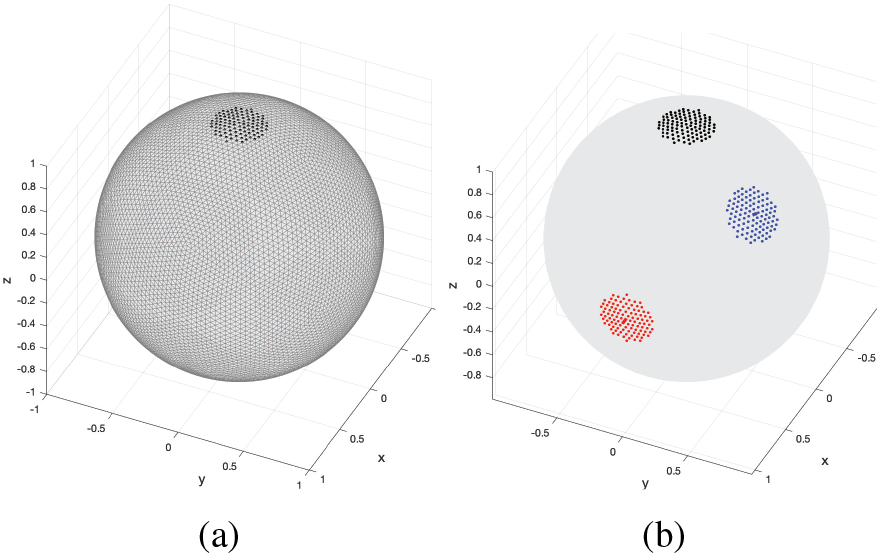
Uniform sampling within solid angles along different orientations. (a) An icosahedron with five levels of subdivision, wherein the subset of its vertices that fall within the solid angle 4*π*/98 around the z-axis direction, marked with black dots, are treated as a template sampling pattern. (b) The template sampling pattern (black) is then rotated towards other neighborhood directions; two directions shown here, in red and blue.

## Appendix D Streamline-based phantom construction

For each subject, 10 thousand streamlines, denoted {*s*_*i*_(*x*) ∈ ℝ^3^}_*i*=1…10000_, were generated through deterministic tractography using the method presented by Yeh et al. (2013), as implemented in DSI Studio. A subset of *S* streamlines from a single subject was randomly selected and used as the basis to produce a phantom. Each streamline *s*_*i*_(*x*) was first voxelized, resulting in a vector **s**_*i*_ containing the indices of the voxels through which it passes. A random source point for the activation was then selected, represented by an indicator vector **d**_*i*_ of the same length as **s**_*i*_, wherein a single element of the vector was set to 1 and the remaining elements were set to 0. An adjacency matrix **A**_*i*_ was then defined, specifying that every voxel in **s**_*i*_ is connected to itself and its neighbors within a 3 ×3 × 3 neighborhood, with equal weights adding up to 1. The diffuse activation pattern, denoted **p**_*i*_, was then synthesized as

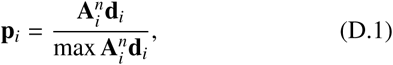

where the exponent *n* is a parameter that controls the extent of spatial spread of the activation. This parameter was arbitrarily set to 250 in the design of all the phantoms used in this work, with the goal of obtaining long and smoothly-decaying spatial activation patterns. Finally, the phantom was constructed by merging the various activation patterns {**p**_*i*_} _*i*=1…*S*_ into a single volume.

## Footnotes

1 https://ida.loni.usc.edu/login.jsp

2 https://www.fil.ion.ucl.ac.uk/spm/software/spm12/

3 http://dsi-studio.labsolver.org

4 https://www.nitrc.org/projects/csaodf-hough

5 https://identifiers.org/neurovault.collection:9494

6 In our default analysis setting, regions outside the WM are masked out of fMRI volumes prior to GSS smoothing. This prevents the introduction of spurious signal, particularly from gray matter, while ensuring an unbiased comparison with DSS. Such considerations are not adhered to when implementing full brain GSS, and these results are therefore provided only for reference. Furthermore, due to the differences in FDR thresholding, there is no expectation of WM detections of either GSS method being a subset of those of the other.

