## Supplementary Results for "Diffusion-Informed Spatial Smoothing of fMRI Data in White Matter Using Spectral Graph Filters"

*Diffusion-informed filters*

Figures S1-S6 illustrate the range of shapes that DSS filters can take by varying their $\tau$ and $\alpha$ parameters. The filters are localized at the three ROIs shown in Figure 5.

The increased angular resolution of the 5-conn graph neighborhood definition allows for narrower and more intricate filter shapes than the 3-conn definition. It also manifests a more straightforward relationship between filter width and parameter $\alpha$. Due to the longer range connections present in the 5-conn definition, filters defined with it tend to be larger than those defined with the 3-conn definition for the same value of $\tau$.

For $\alpha$ values of 0.5, the diffusion-adaptive properties of graph filters become negated, resulting in isotropic filters. Reducing $\alpha$ below 0.5 has increasingly negligible effects on the resulting filters, given the weak nature of the connections it introduces.

*ROC analysis*

Figure S7(a) shows an overview of the ROC performance of DSS and GSS on the single-volume streamline-based phantoms. Given the number of phantoms, noise realizations, and settings used in our tests, presenting the individual ROC curves would be unfeasible. Instead, we present ensemble ROC curves, computed by averaging the 950 individual ROC curves for each given setting. Given that the individual ROC curves were defined by non-uniformly spaced sets of FPRs and TPRs, each individual ROC curve was first interpolated to obtain a continuous representation, and then sampled at 300 TPR levels uniformly distributed between 0 and 1. Lastly, an average FPR was computed for each TPR level. These results show that DSS outperforms GSS in terms of both sensitivity and specificity across the range of filter sizes and settings tested.

Figures S7(b) and (c) present results of the denoising performance of DSS and GSS on single-volume streamline-based phantoms with varying SNR. The ground truth activations in streamline-based phantoms take values in the $[0, 1]$ range. Secondary phantoms were created from these, with values in the $[\theta, 1]$ range. By discarding activation values below $\theta$, while keeping the same noise standard deviation of 1, the SNR of the phantoms was increased. These results show that DSS outperforms GSS over a range of SNR values.

*Single-subject activation mapping results*

Figures S11 and S12 present comparisons of single-subject activation mapping results obtained with GSS and DSS. Despite the presence of deep WM activations, both methods can also manifest activations close to the GM boundary. These vary considerably in number and extent, and while they may be attributed to partial volume effects. However, the full brain activation maps reveal that there is not complete parity between active GM regions and detections by GSS and DSS along the WM border.

The following is a breakdown of notable observations from each activation mapping comparison:

- In Figure S11(a), DSS detects two parallel streamline-shaped activations for $\tau$ values of 3 and over (orange arrow). When using GSS, the second activation is either not detected or combined with the first. Notably, the GSS activation is elongated for small filters of 1 and 2 mm FWHM, but becomes round for larger filter sizes, suggesting loss of spatial specificity.
- In Figure S11(b), DSS detects two activations which, while somewhat overlapping, can be identified as distinct streamline-shaped activations (orange arrow). GSS is generally capable of detecting both activations, but they become merged in perpendicular to the streamline orientation, preventing their identification as parallel axonal bundles.
- Figure S12(a), presents an example where both GSS and DSS are capable of successfully resolving two parallel streamline-shaped activations (orange arrow). However, while larger GSS filter sizes eventually merge both activations into a single large blob, their shape and extent remains fairly constant across DSS filter sizes.
- Figure S12(b) presents activations on a sagittal slice across the corpus callosum (orange and yellow arrows). Given that axonal bundles are oriented in perpendicular to the image plane, the size of DSS activations stays relatively constant across filter sizes, allowing precise localization of callosal activations. On the other hand, with GSS, the size of one of these activations increases in proportion to filter size (orange arrow), as the data are smoothed isotropically in 3-D, while another is present only for a narrow range of filter sizes (yellow arrow).

*Group activation mapping results*

Figures S14-S16 present comparisons of group activation mapping results obtained with GSS and DSS. The following is a breakdown of notable observations from each activation mapping comparison:

- In Figure S14(a), DSS produces two distinct streamline-shaped activations across all of the tested filter sizes (orange and yellow arrows). Only one of these activations is also produced by GSS (orange arrow), and only for the largest filter sizes tested.
- Figure S14(b) presents a similar case, where DSS detects a distinct streamline-shaped activation across all of the tested filters sizes (orange arrow), while GSS displays a much smaller activation, and only for the largest of filter sizes tested.
- Figure S15(a) present several phenomena of interest. Firstly, DSS detects a slender streamline-shaped activation across all filters sizes (orange arrow), while GSS detections are at the same location are smaller and rounder. Secondly, large DSS filters detect two adjacent streamline-shaped activations that remain distinct (yellow arrow), while GSS only detects one of them, as a round shape, with the largest filter size tested.

- Figure S15(b) shows an activation in the corpus callosum which is detected by DSS across all filter sizes (orange arrow), but not by GSS. Notably, the t-value produced by GSS at this region is inversely proportional to the size of the filter used, illustrating the negative effects of isotropic smoothing for the detection of small, intricate activation patterns.

- Figure S16(a) GSS detects several elongated activation patterns (orange arrows), but these have greater spatial extent and are detected more consistently with DSS, which also detects an additional streamline-shaped activation (yellow arrow) that is almost completely undetected by GSS.

- In Figure S16(b), DSS produces a single streamline-shaped activation pattern (orange arrow), while GSS does not show an activation at the same region.

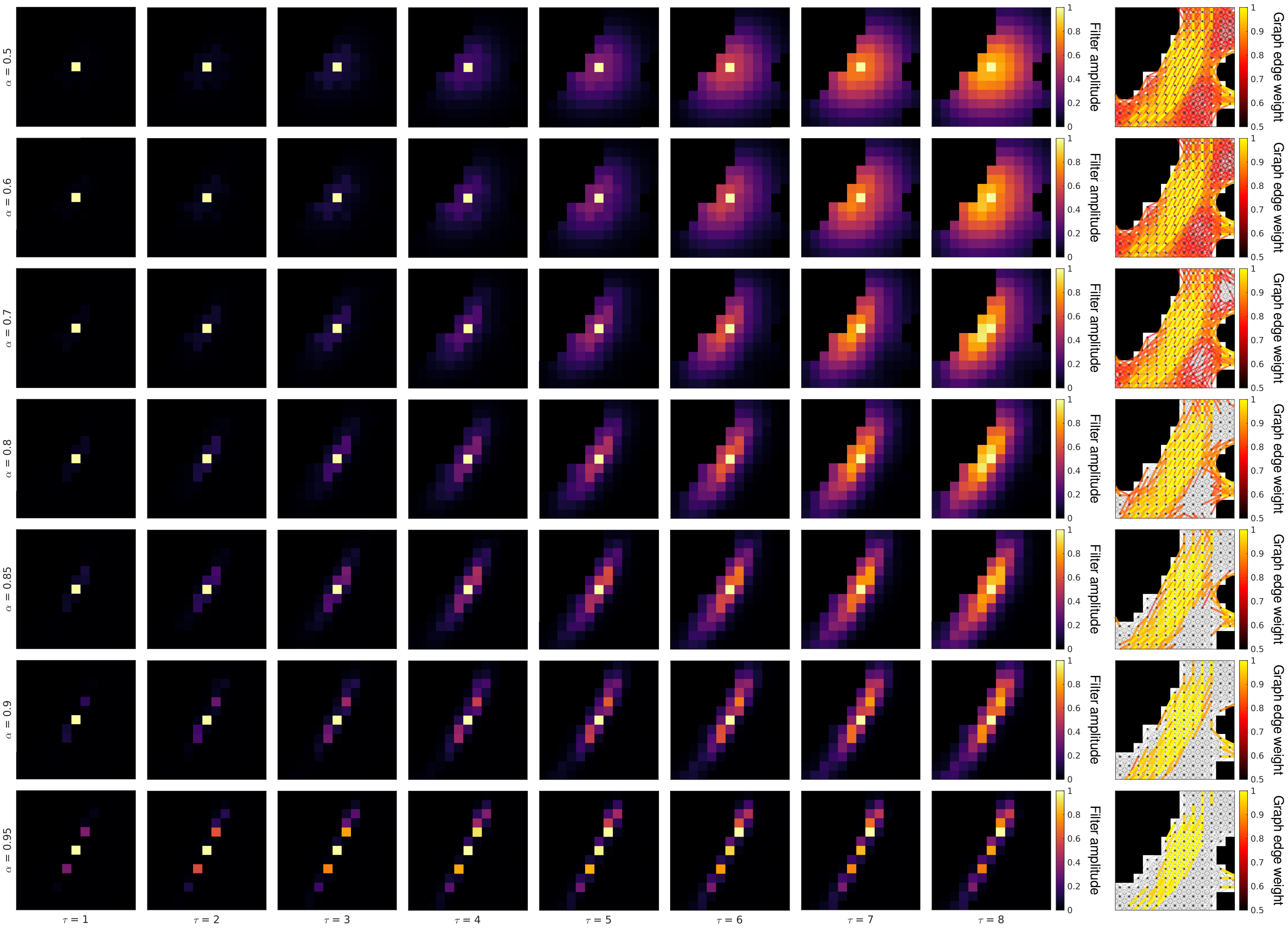


Figure S1: Effects of parameters $\tau$ and $\alpha$ on the shape of DSS filters when realized at the center of red ROI shown in Figure 5. Graph parameters: 5-conn neighborhood, $\beta = 50$. Filters are shown normalized to the $[0, 1]$ range.

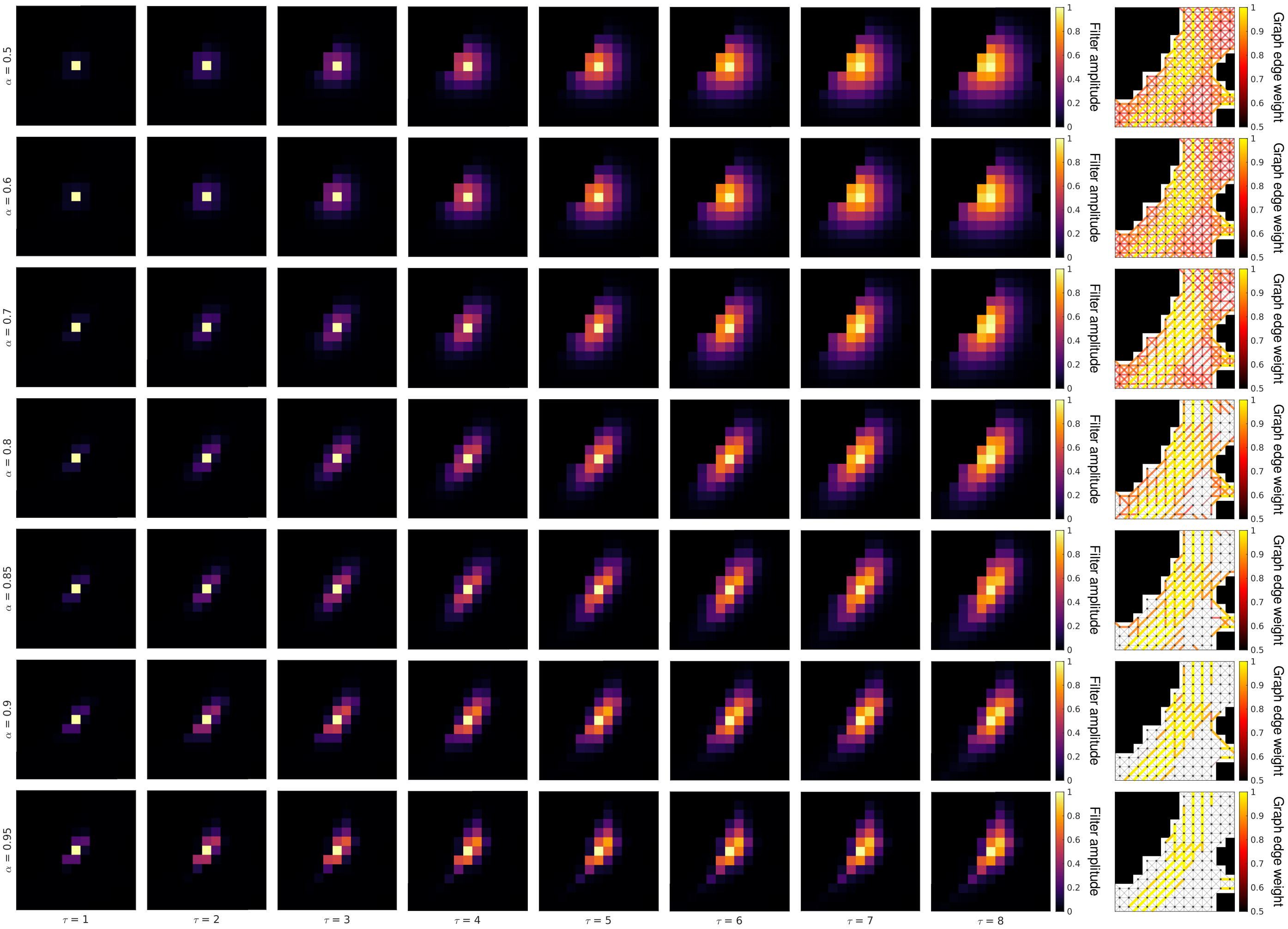


Figure S2: Effects of parameters $\tau$ and $\alpha$ on the shape of DSS filters when realized at the center of the red ROI shown in Figure 5. Graph parameters: 3-conn neighborhood, $\beta = 50$. Filters are shown normalized to the $[0, 1]$ range.

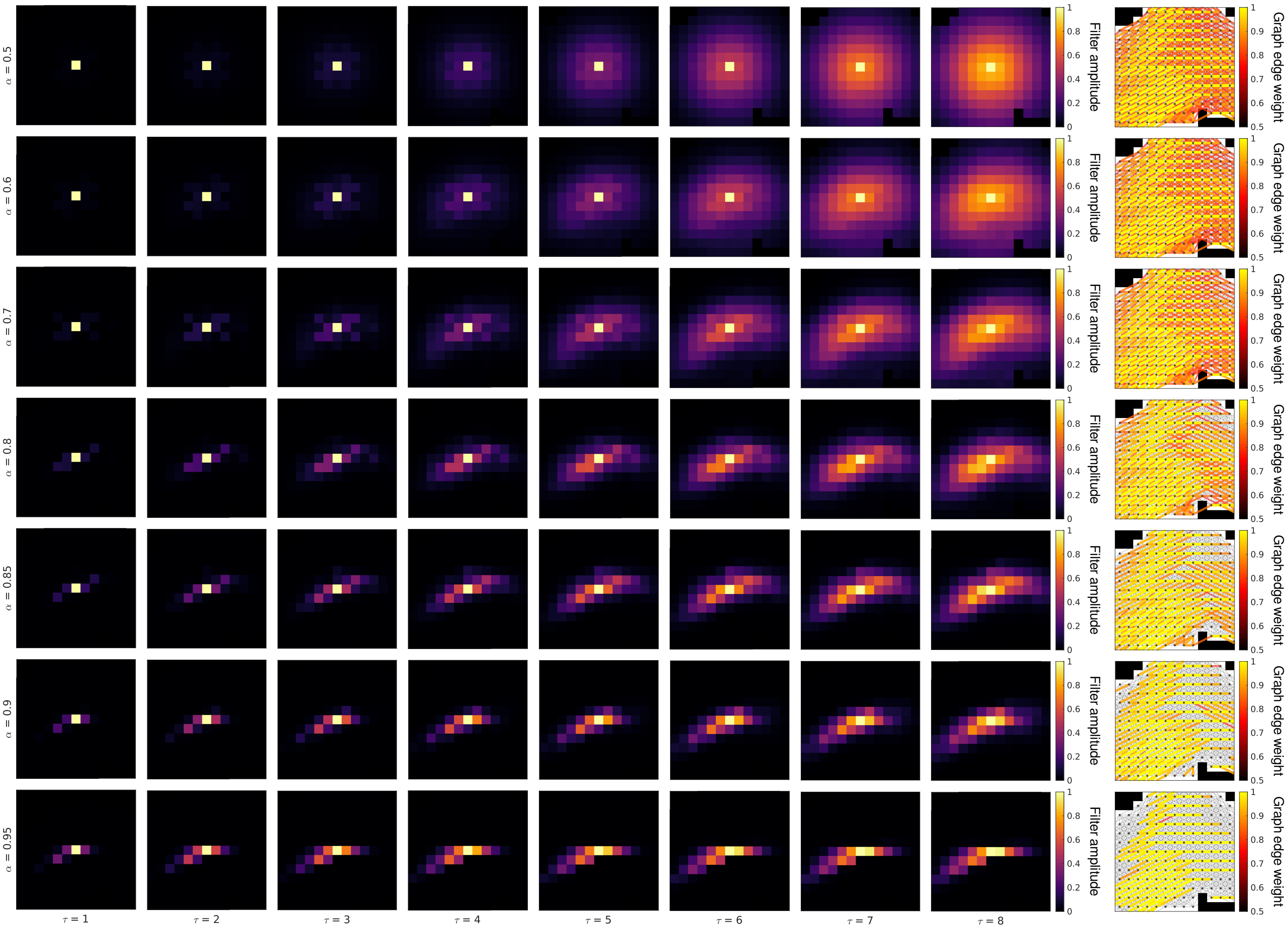


Figure S3: Effects of parameters $\tau$ and $\alpha$ on the shape of DSS filters when realized at the center of the blue ROI shown in Figure 5. Graph parameters: 5-conn neighborhood, $\beta = 50$. Filters are shown normalized to the $[0, 1]$ range.

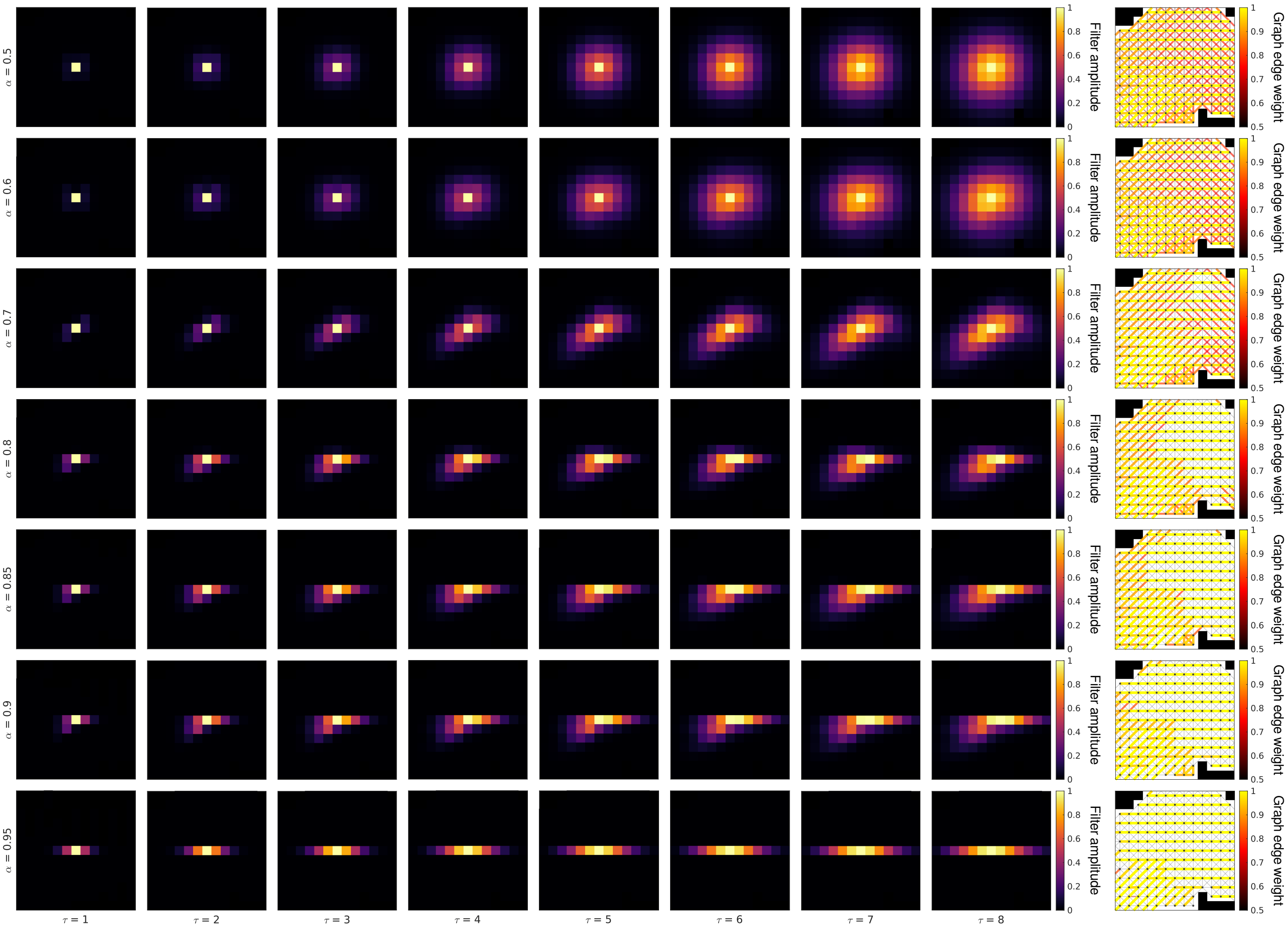


Figure S4: Effects of parameters $\tau$ and $\alpha$ on the shape of DSS filters when realized at the center of the blue ROI shown in Figure 5. Graph parameters: 3-conn neighborhood, $\beta = 50$. Filters are shown normalized to the $[0, 1]$ range.

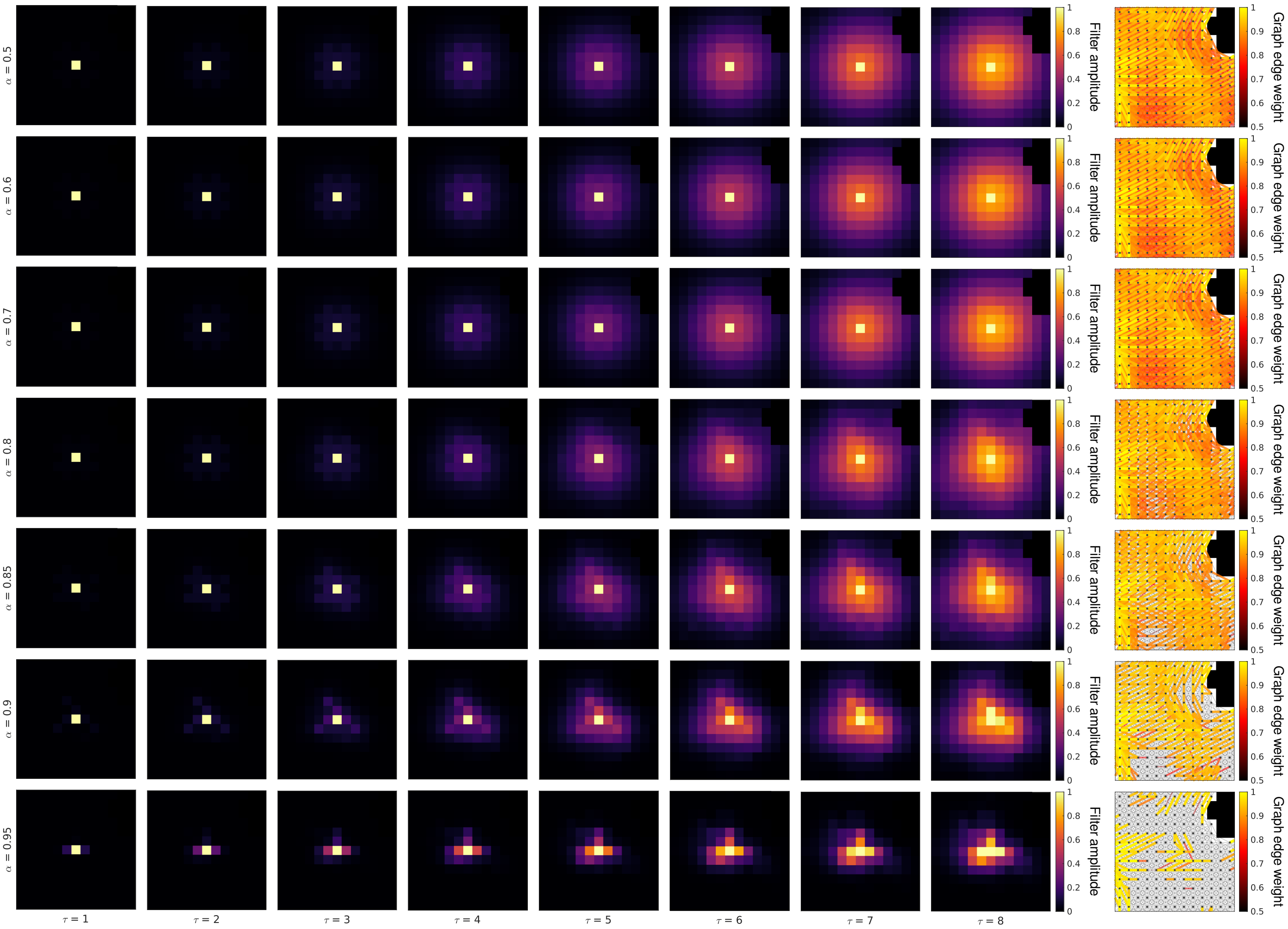


Figure S5: Effects of parameters $\tau$ and $\alpha$ on the shape of DSS filters when realized at the center of the yellow ROI shown in Figure 5. Graph parameters: 5-conn neighborhood, $\beta = 50$. Filters are shown normalized to the [0, 1] range.

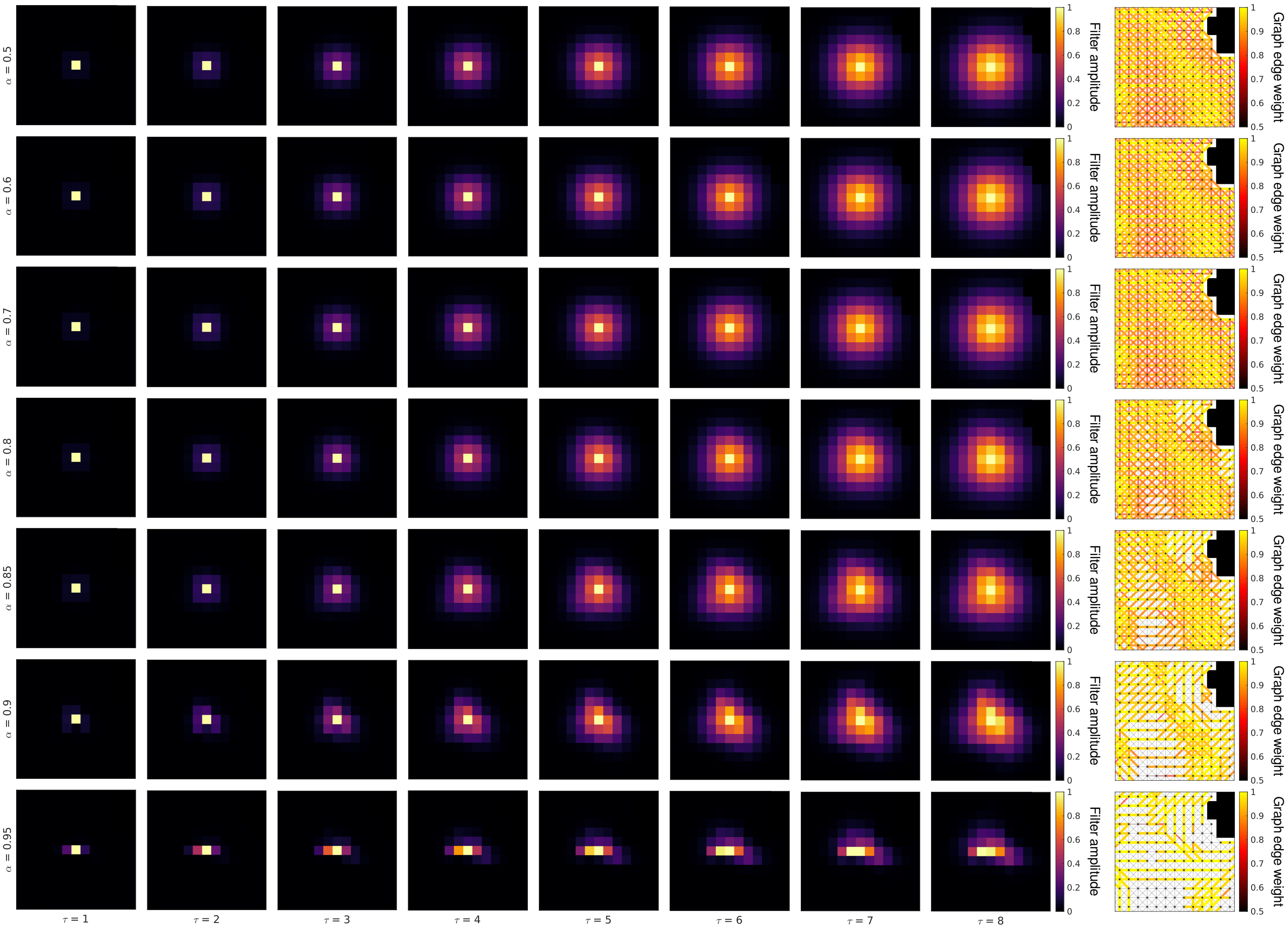


Figure S6: Effects of parameters $\tau$ and $\alpha$ on the shape of DSS filters when realized at the center of the yellow ROI shown in Figure 5. Graph parameters: 3-conn neighborhood, $\beta = 50$. Filters are shown normalized to the $[0, 1]$ range.

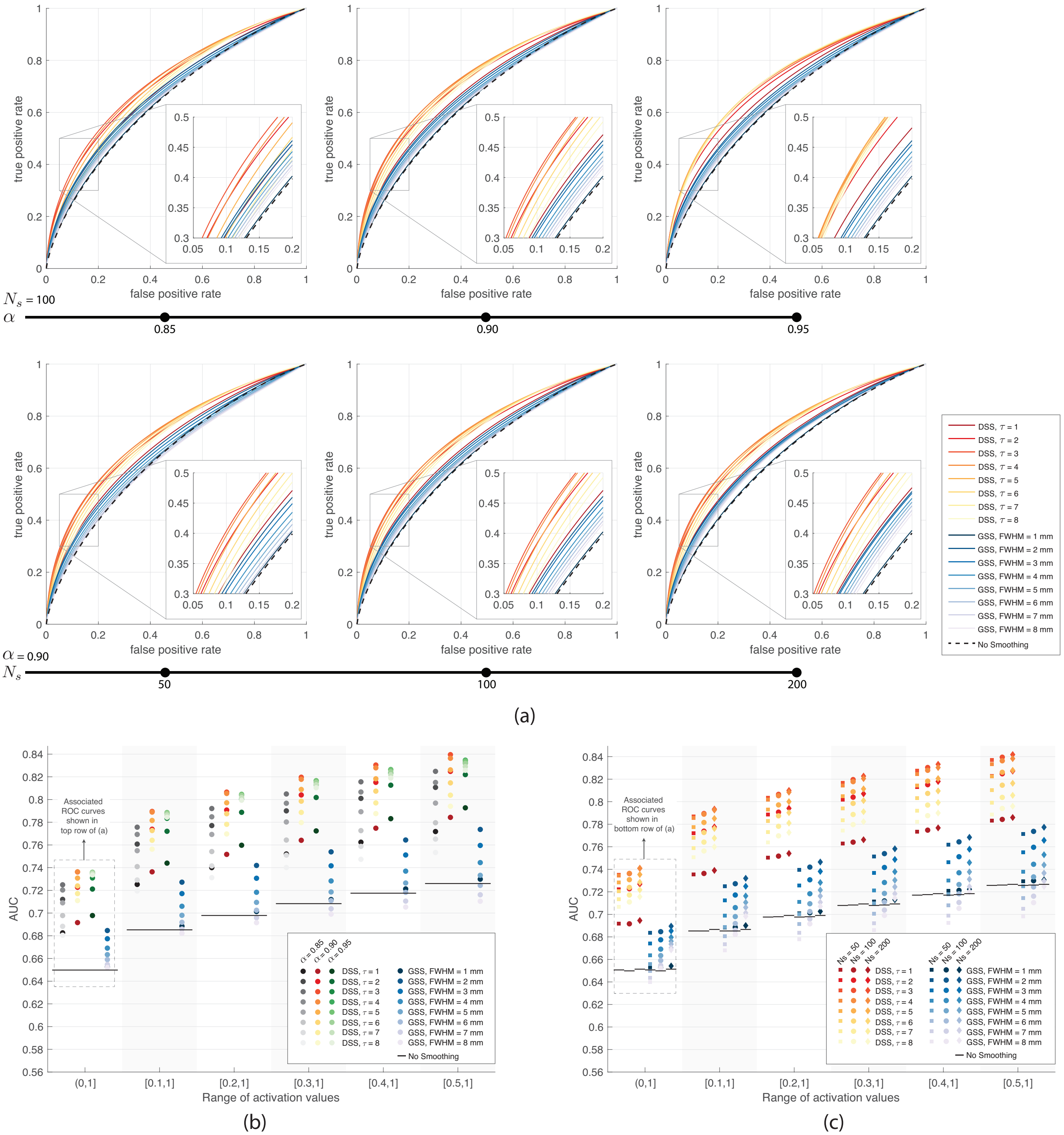


Figure S7: (a) ROC analysis of denoising performance by GSS and DSS on streamline-based phantoms. The three panels on the top compare the performance of DSS for three values of $\alpha$ on the streamline-based phantoms with $N_s$ = 100; the ROC curves associated to GSS are identical for these three settings. The three panels on the bottom compare the performance of DSS and GSS on streamline-based phantoms with $N_s$ = 50, 100 and 200; a fixed value of $\alpha$ = 0.9 is used for DSS in these three settings. The two panels in the central column are identical. (b)-(c) AUCs of ensemble ROC curves obtained on spatially smoothed single-volume streamline-based phantoms with varying SNR. (b) Performance of GSS is compared with that of DSS with a range of $\alpha$ values, on phantoms with $N_s$ = 100. (c) Performance of GSS is compared with that of DSS with a fixed value of $\alpha$ = 0.9, on phantoms with $N_s$ = 50, 100 and 200.

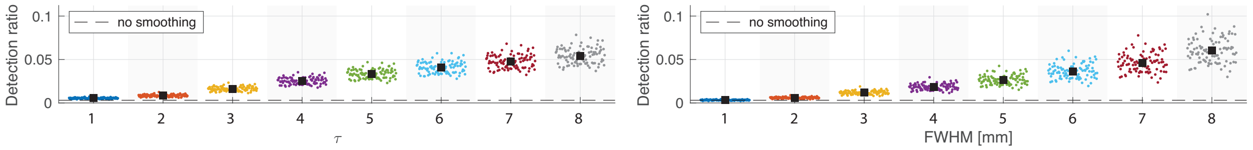


(a)

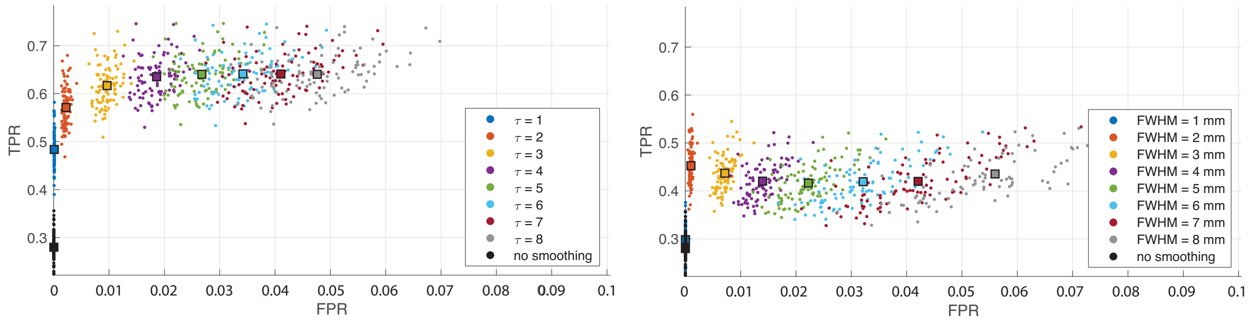


(b)

Figure S8: (a) Fraction of voxels within WM mask detected as being significant using DSS (left) and GSS (right) across the 95 time-series streamline-based phantoms with 50 streamlines. Significant voxels were determined after FDR correction at 5%. (b) The corresponding TPR and FPR for DSS and GSS are shown in the left and right, respectively. Each scatter dot corresponds to one phantom, whereas ■ in the top row and color-filled □ in the bottom row show the average values across the 95 phantoms.

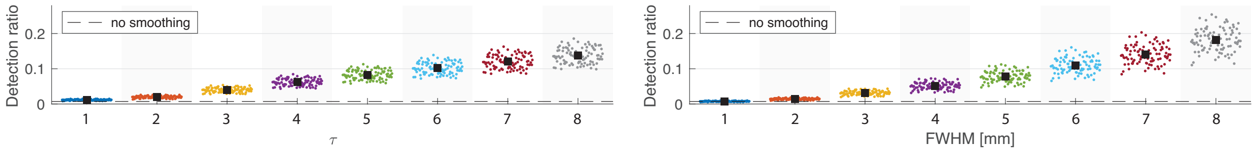


(a)

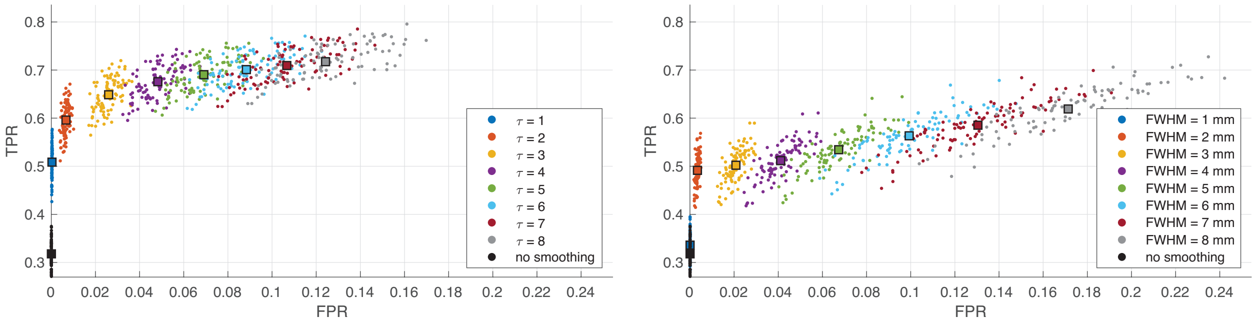


(b)

Figure S9: Same as Figure S8, but on time-series streamline-based phantoms with 100 streamlines.

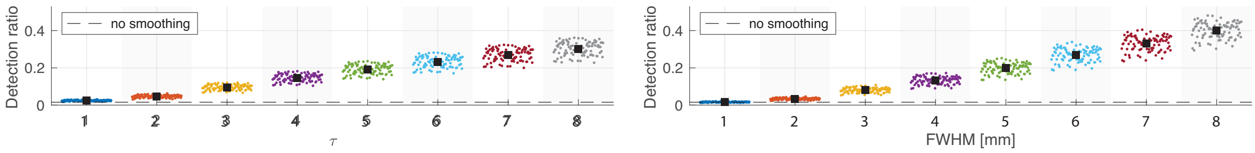


(a)

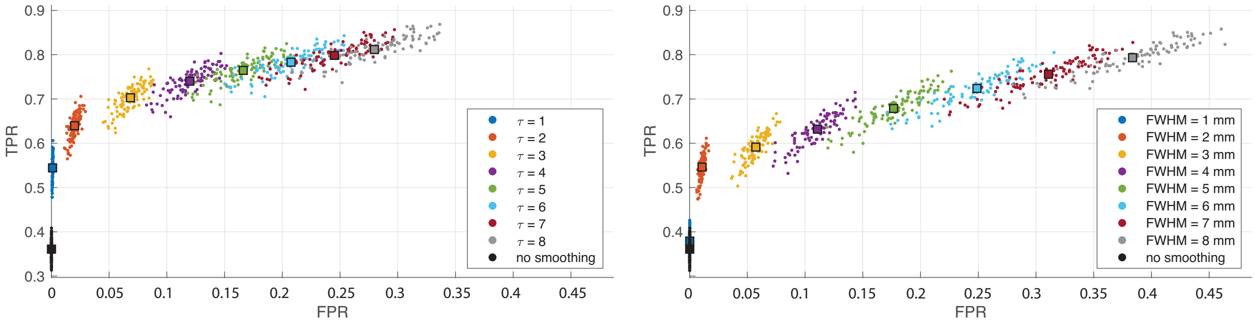


(b)

Figure S10: Same as Figure S8, but on time-series streamline-based phantoms with 200 streamlines.

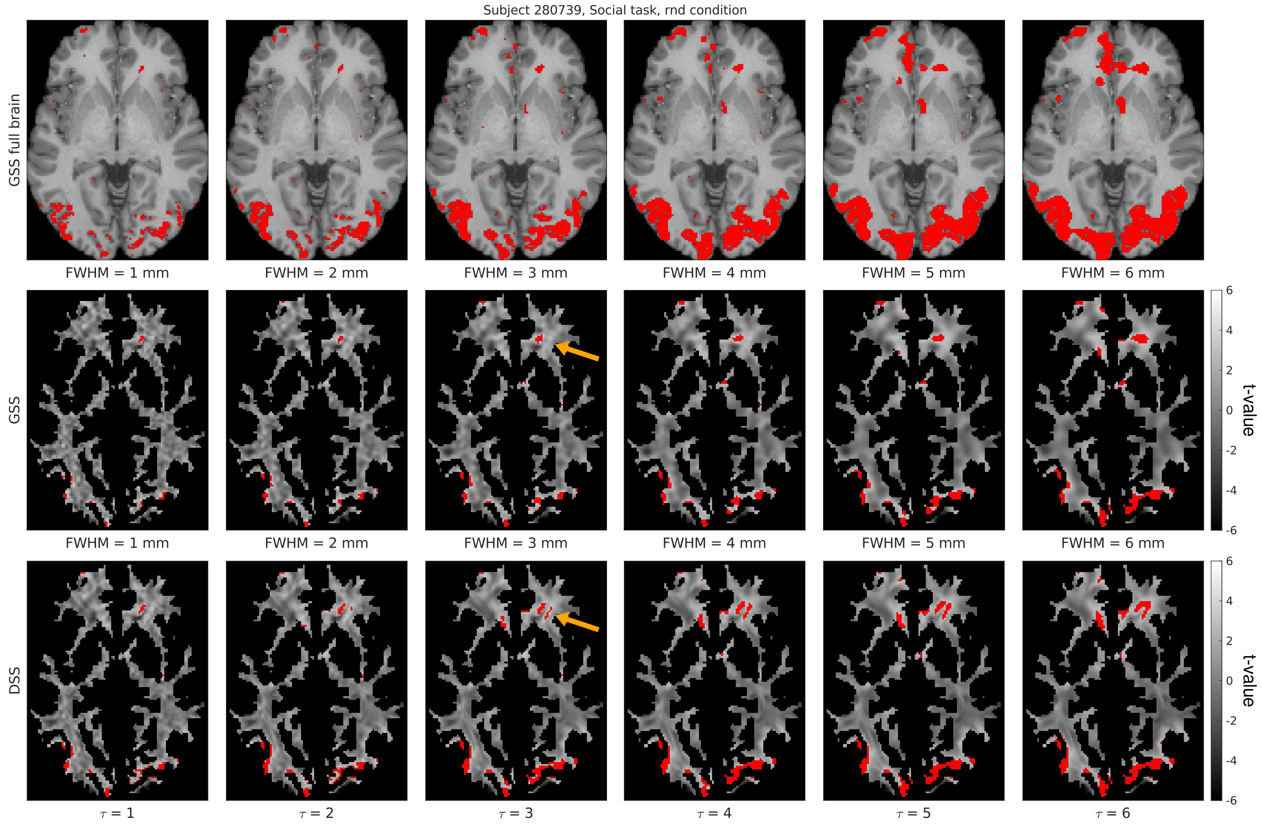


(a)

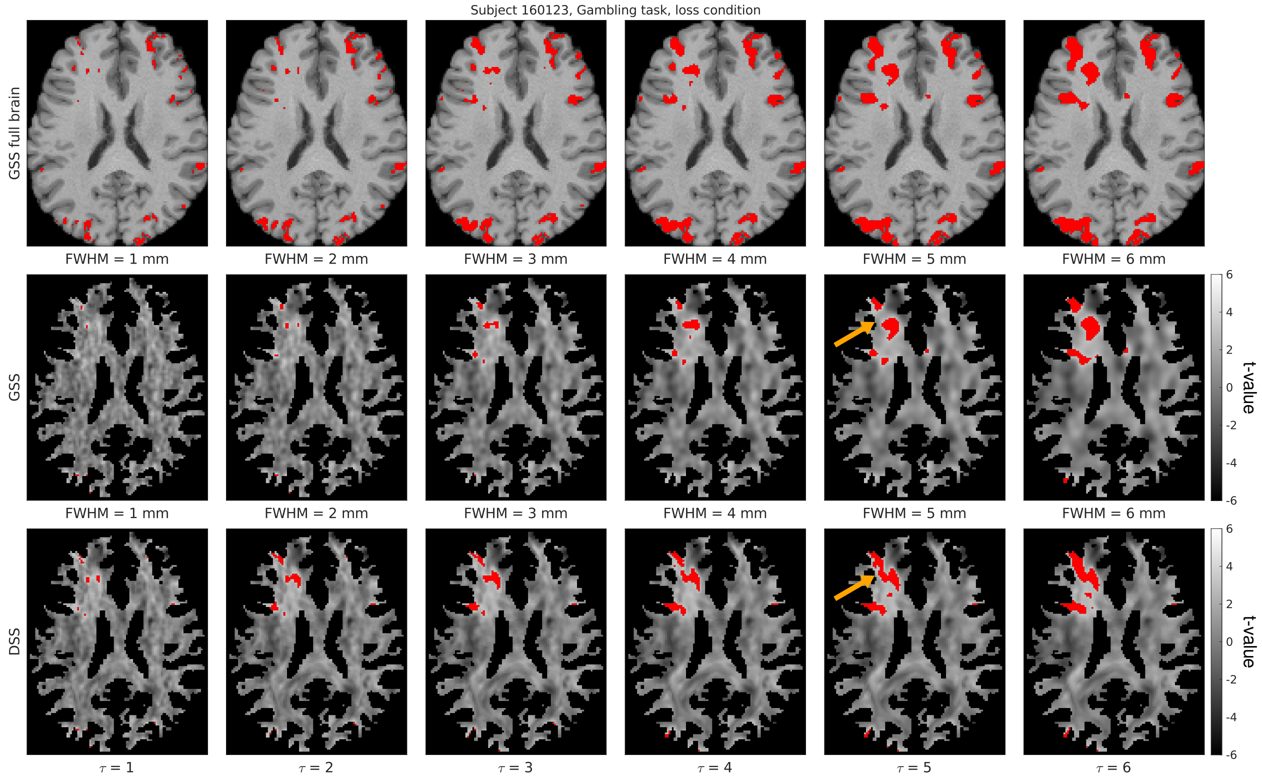


(b)

Figure S11: Comparison of representative single-subject activation mapping results generated with GSS and DSS, with t-maps shown in grayscale and detections overlaid in red (FDR-corrected at 5%). Full-brain activation maps are also shown for reference, overlaid on the subject's T1w image.

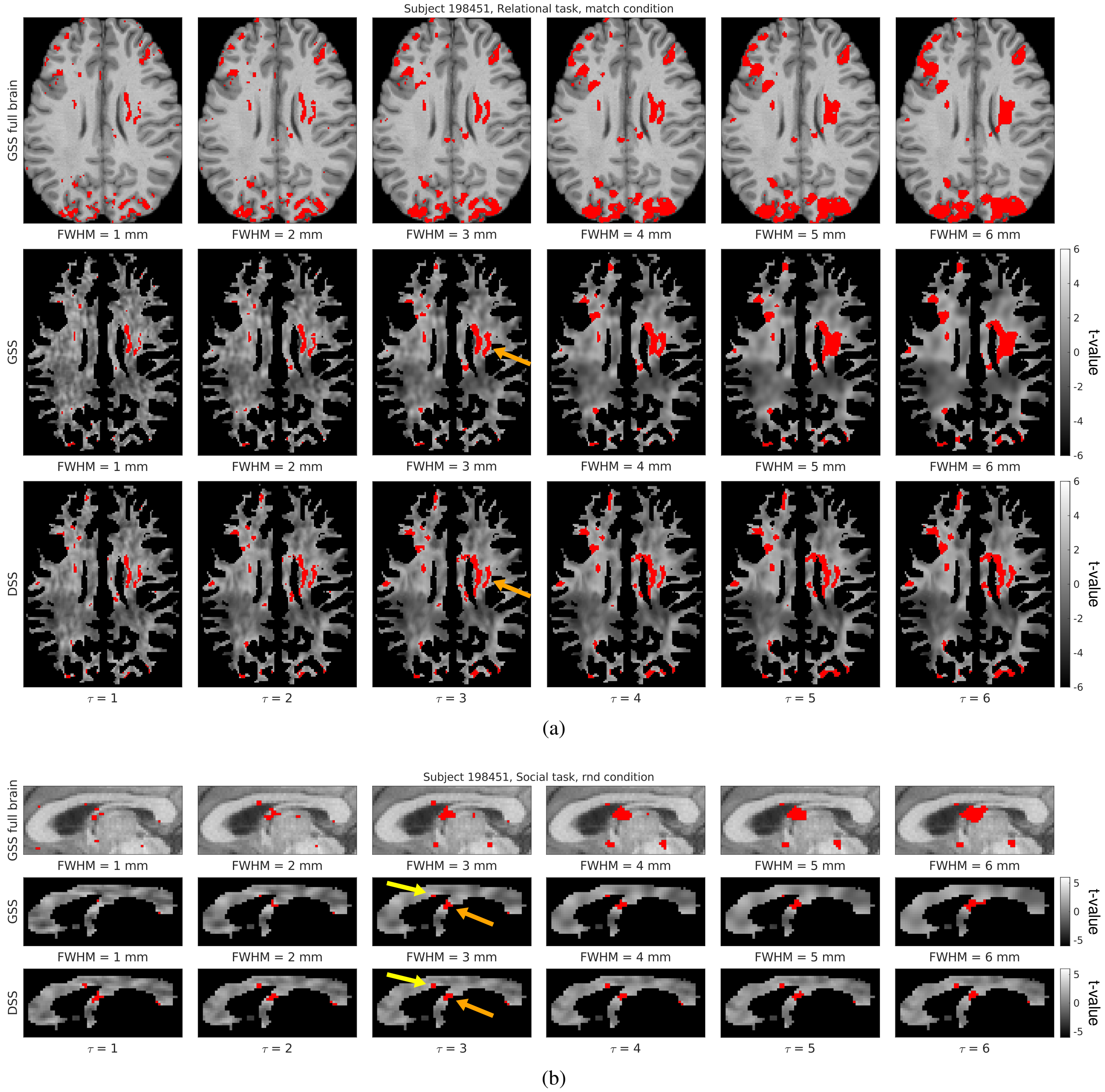


Figure S12: Comparison of representative single-subject activation mapping results generated with GSS and DSS, with t-maps shown in grayscale and detections overlaid in red (FDR-corrected at 5%). Full-brain activation maps are also shown for reference, overlaid on the subject's T1w image.

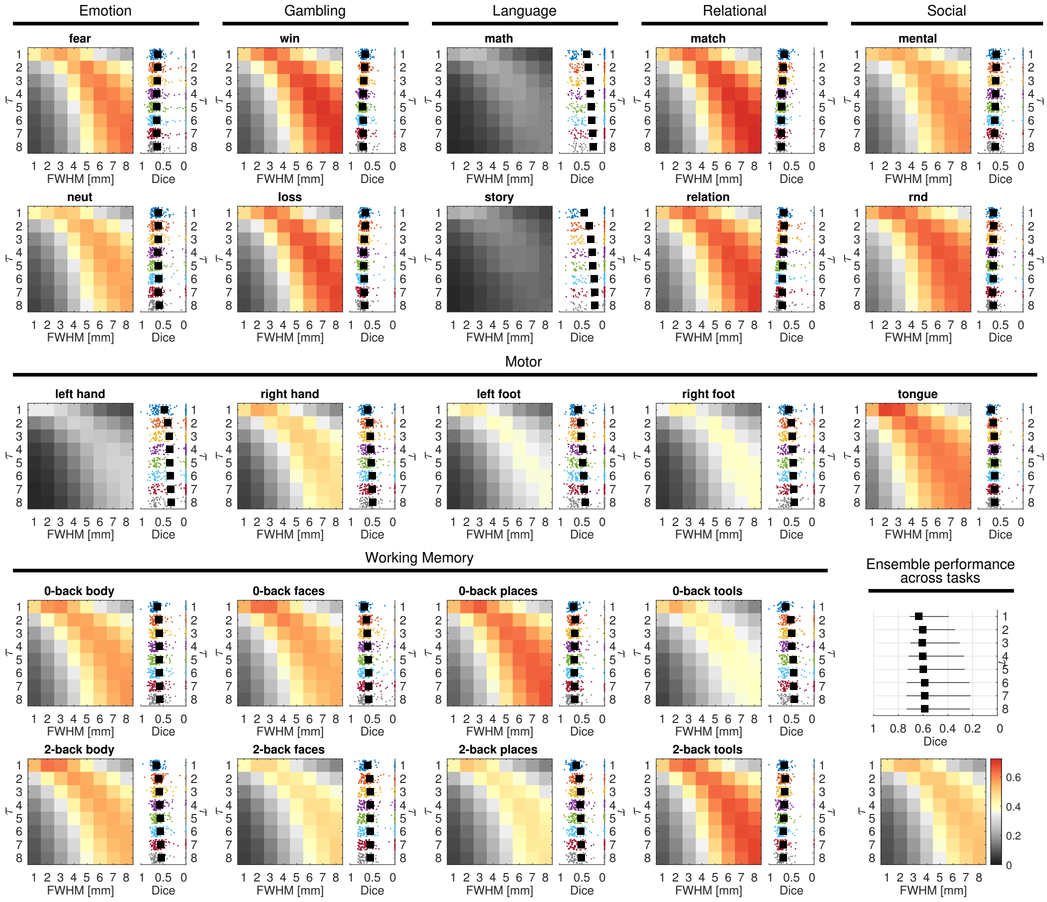


Figure S13: Dice similarity between detection maps generated with DSS and GSS across all 23 experimental conditions of the seven HCP functional tasks. See Figure 12 and its caption for a full description.

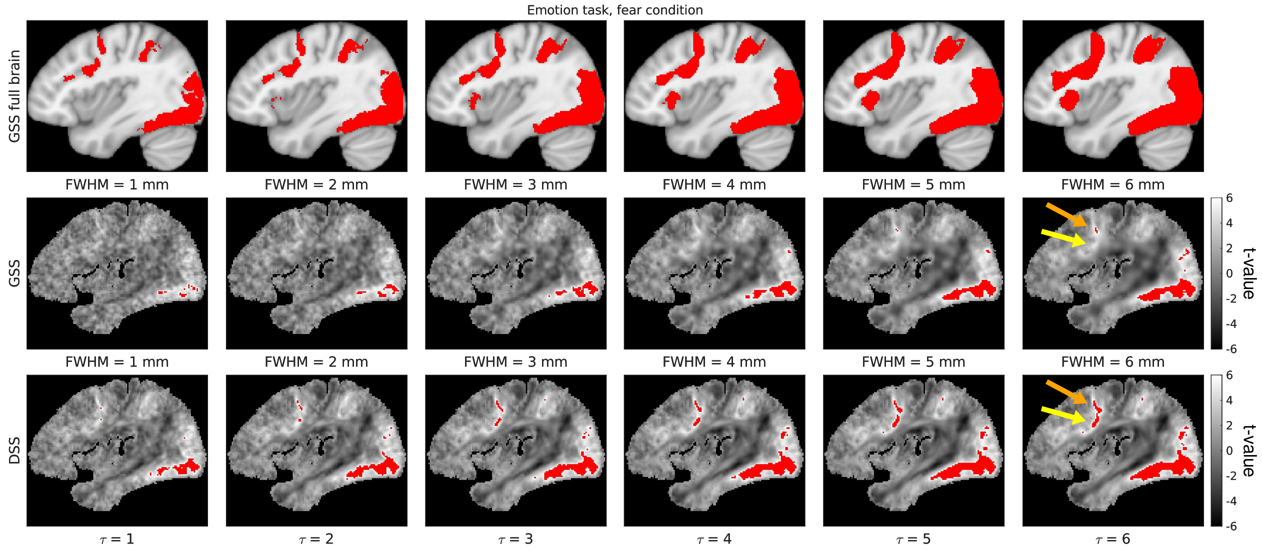


(a)

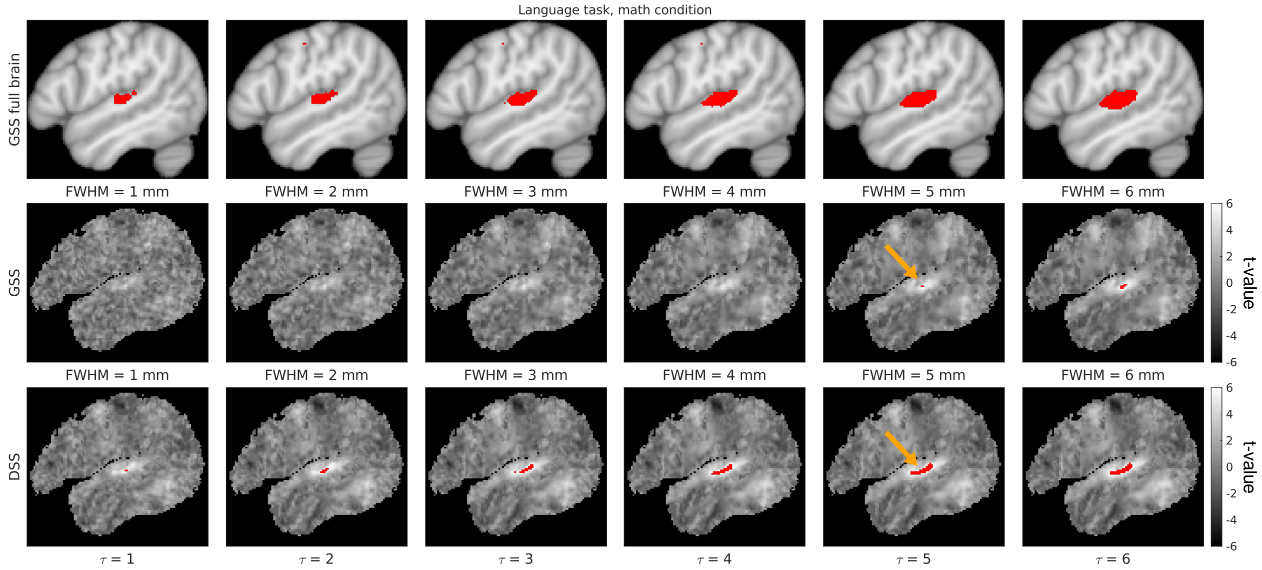


(b)

Figure S14: Comparison of representative group activation mapping results generated with GSS and DSS, with t-maps shown in grayscale and detections overlaid in red (Bonferroni-corrected at 5%). Full-brain activation maps are also shown for reference, overlaid on the MNI152 T1w template image.

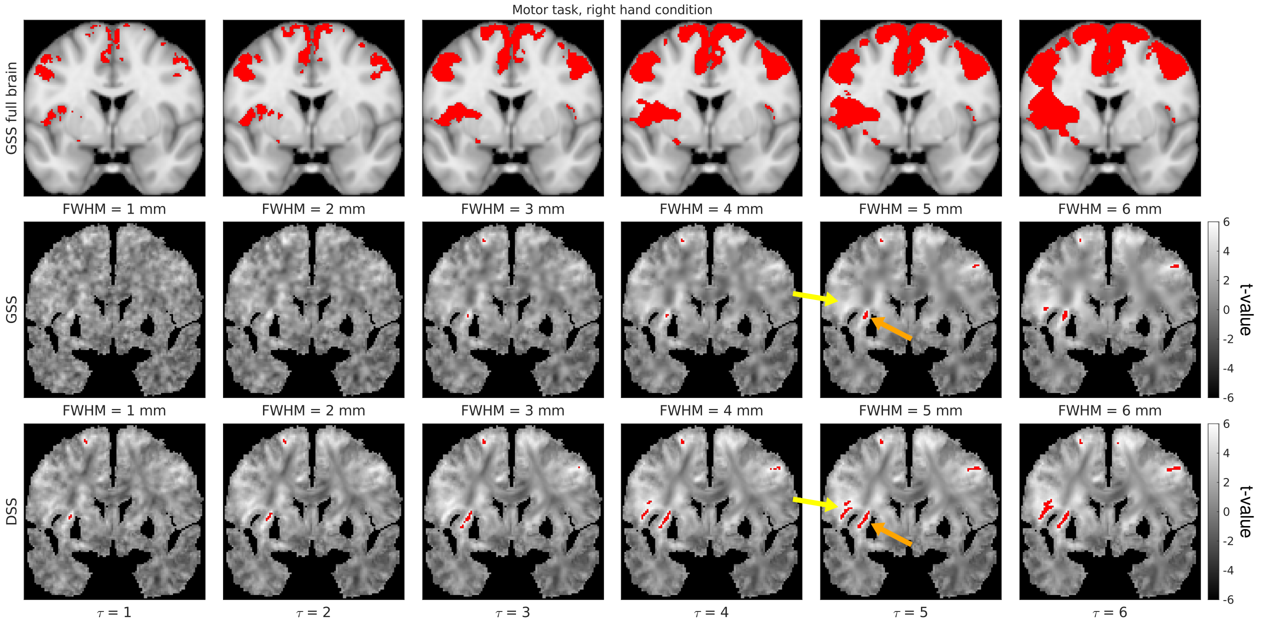


(a)

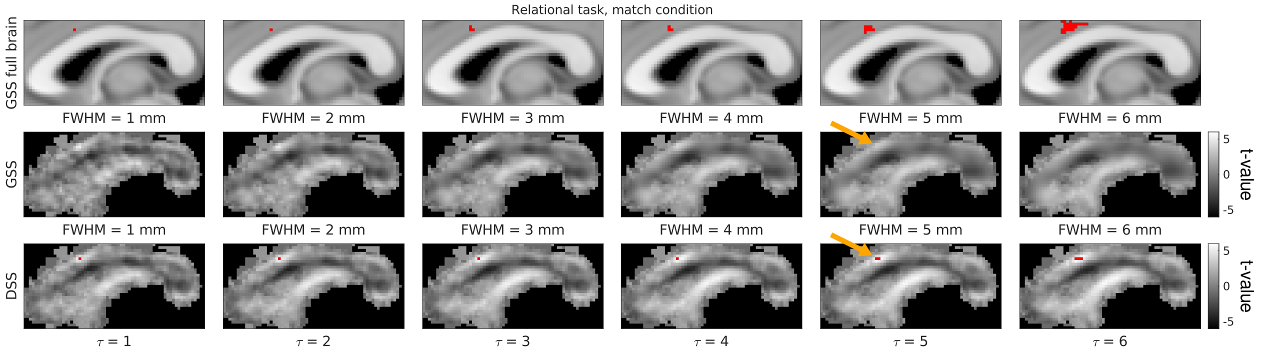


(b)

Figure S15: Comparison of representative group activation mapping results generated with GSS and DSS, with t-maps shown in grayscale and detections overlaid in red (Bonferroni-corrected at 5%). Full-brain activation maps are also shown for reference, overlaid on the MNI152 T1w template image.

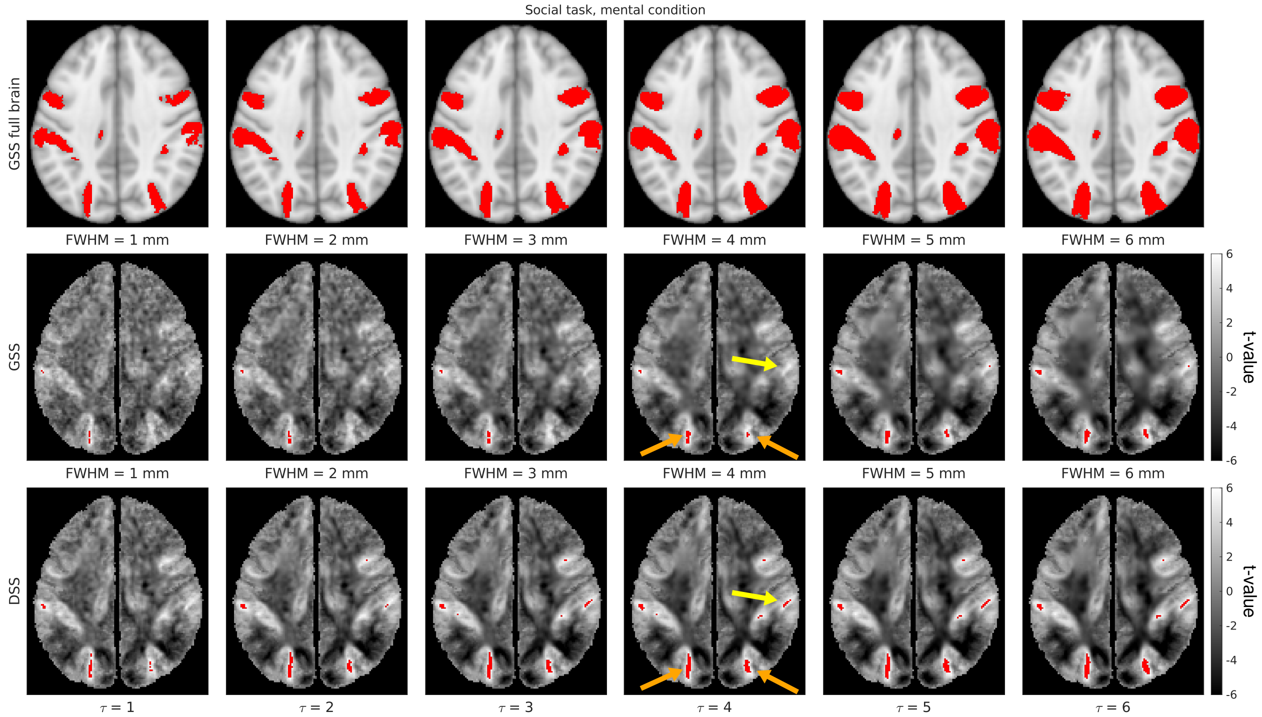


(a)

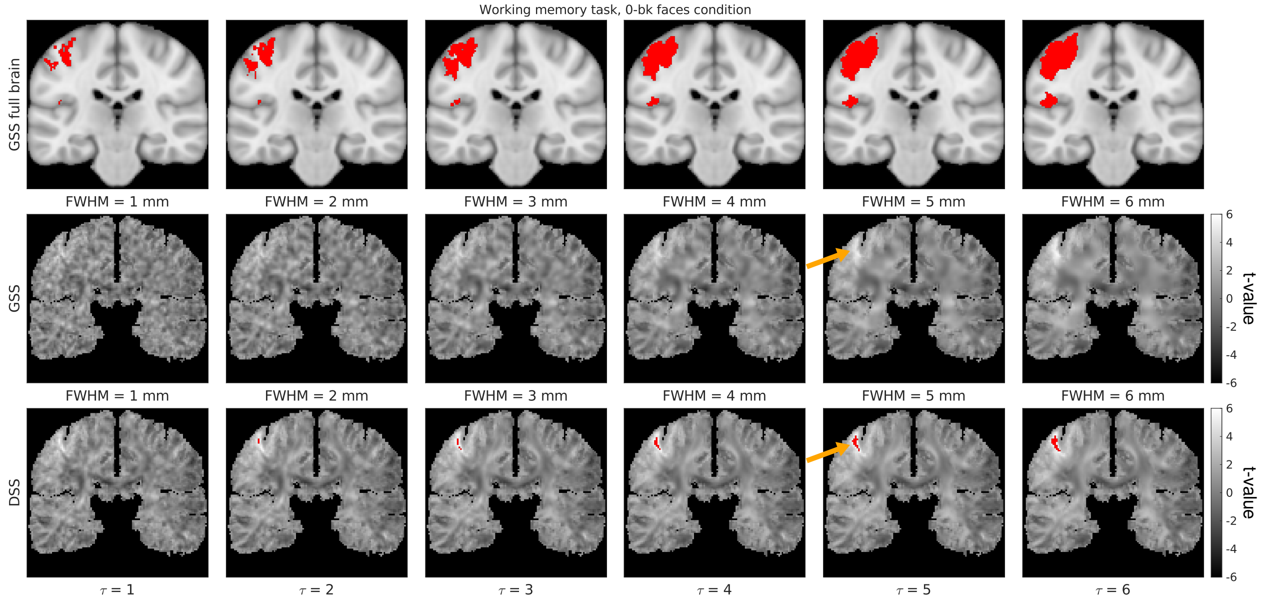


(b)

Figure S16: Comparison of representative group activation mapping results generated with GSS and DSS, with t-maps shown in grayscale and detections overlaid in red (Bonferroni-corrected at 5%). Full-brain activation maps are also shown for reference, overlaid on the MNI152 T1w template image.

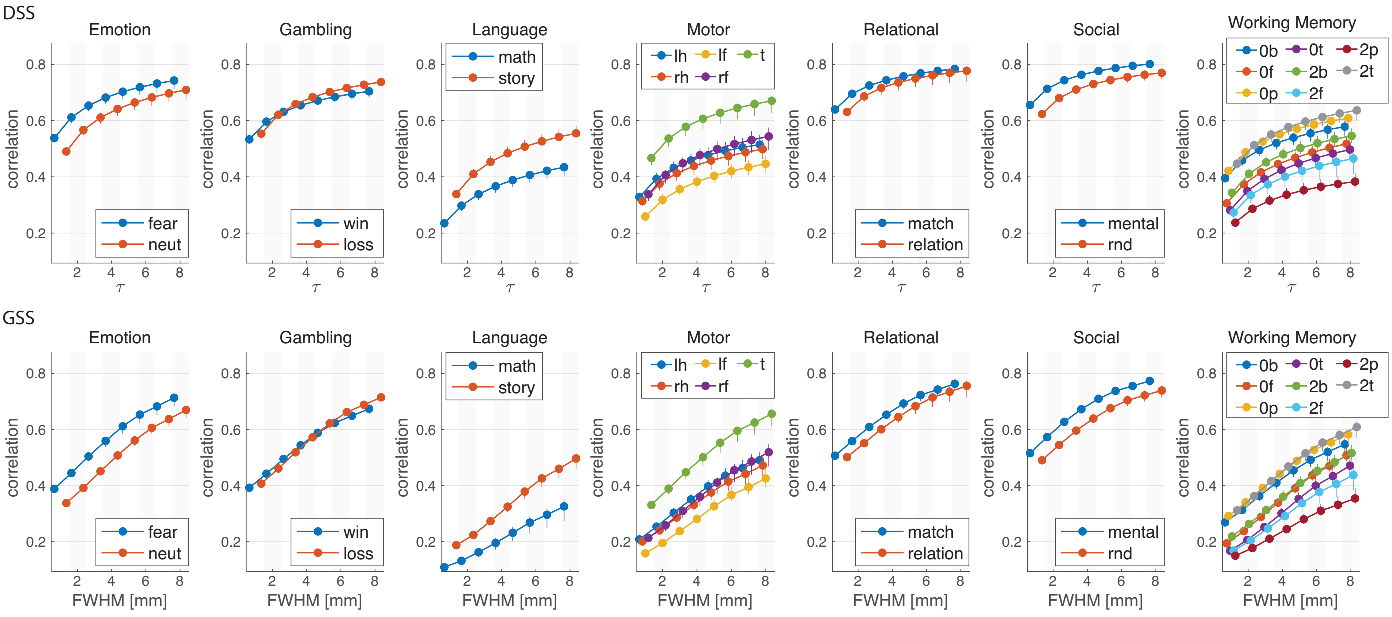


(a)

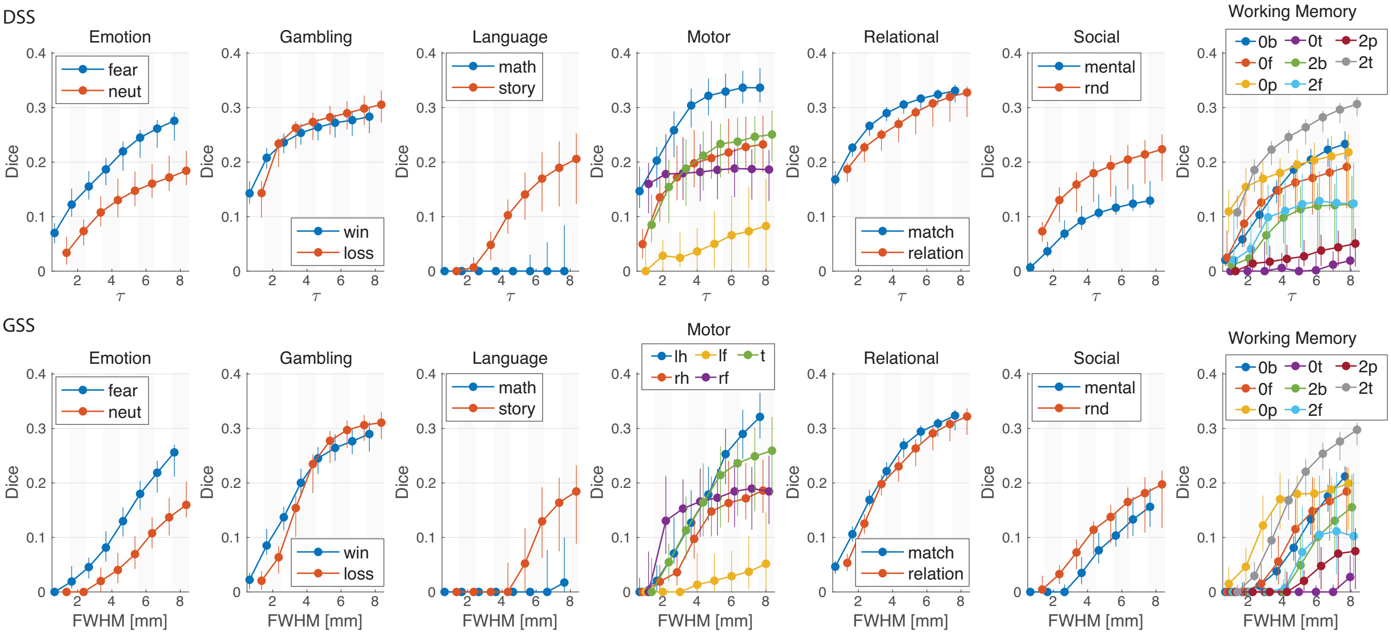


(b)

Figure S17: Results of Monte Carlo test-retest analysis for all experimental conditions. Subjects were repeatedly divided into two groups and subjected to group analysis, and the resulting statistical maps were compared. (a) Correlation between t-maps of both groups. (b) Dice similarity between activation maps of both groups. The markers show the median value across 30 experiments, whereas the whiskers represent 5 – 95% percentiles. Abbreviations: lh: left hand, rh: right hand, lf: left foot, rf: right foot, t: tongue, 0b/f/p/t: 0-back body/faces/places/tools, 2b/f/p/t: 2-back body/faces/places/tools.
